# Integrative analysis of the shikonin metabolic network identifies new gene connections and reveals evolutionary insight into shikonin biosynthesis

**DOI:** 10.1101/2021.06.30.450579

**Authors:** Thiti Suttiyut, Robert P. Auber, Manoj Ghaste, Cade N. Kane, Scott A. M. McAdam, Jennifer H. Wisecaver, Joshua R. Widhalm

## Abstract

Plant specialized 1,4-naphthoquinones present a remarkable case of convergent evolution. Species across multiple discrete orders of vascular plants produce diverse 1,4-naphthoquinones via one of several pathways using different metabolic precursors. Evolution of these pathways was preceded by events of metabolic innovation and many appear to share connections with biosynthesis of photosynthetic or respiratory quinones. Here, we sought to shed light on the metabolic connections linking shikonin biosynthesis with its precursor pathways and on the origins of shiknoin metabolic genes. Downregulation of *Lithospermum erythrorhizon* geranyl diphosphate synthase (LeGPPS), recently shown to have been recruited from a cytoplasmic farnesyl diphosphate synthase (FPPS), resulted in reduced shikonin production and a decrease in expression of mevalonic acid and phenylpropanoid pathway genes. Next, we used *LeGPPS* and other known shikonin pathway genes to build a coexpression network model for identifying new gene connections to shikonin metabolism. Integrative in silico analyses of network genes revealed candidates for biochemical steps in the shikonin pathway arising from Boraginales-specific gene family expansion. Multiple genes in the shikonin coexpression network were also discovered to have originated from duplication of ubiquinone pathway genes. Taken together, our study provides evidence for transcriptional crosstalk between shikonin biosynthesis and its precursor pathways, identifies several shikonin pathway gene candidates and their evolutionary histories, and establishes additional evolutionary links between shikonin and ubiquinone metabolism. Moreover, we demonstrate that global coexpression analysis using limited transcriptomic data obtained from targeted experiments is effective for identifying gene connections within a defined metabolic network.

## Introduction

The shikonins are a group of red-pigmented naphthoquinones produced in the root periderm of many members of Boraginaceae^1, 2^. They include shikonin (Fig. 1), its enantiomer alkannin, and several shikonin/alkannin derivatives that are excreted into the rhizosphere, where they function in defense, mediate plant-microbe interactions, and/or elicit allelopathic effects on other plants. For example, the invasion success of Paterson’s curse (*Echium plantagineum*) in southeast Australia is attributed, at least in part, to the synthesis and release of shikonins^3^. Shikonins are also the bioactive compounds responsible for the various pharmacological properties of medicinal plants like red gromwell (*Lithospermum erythrorhizon*)^4^ and have emerged as scaffolds for semi-synthesis of novel cancer therapeutics^5^.

**Fig. 1.**
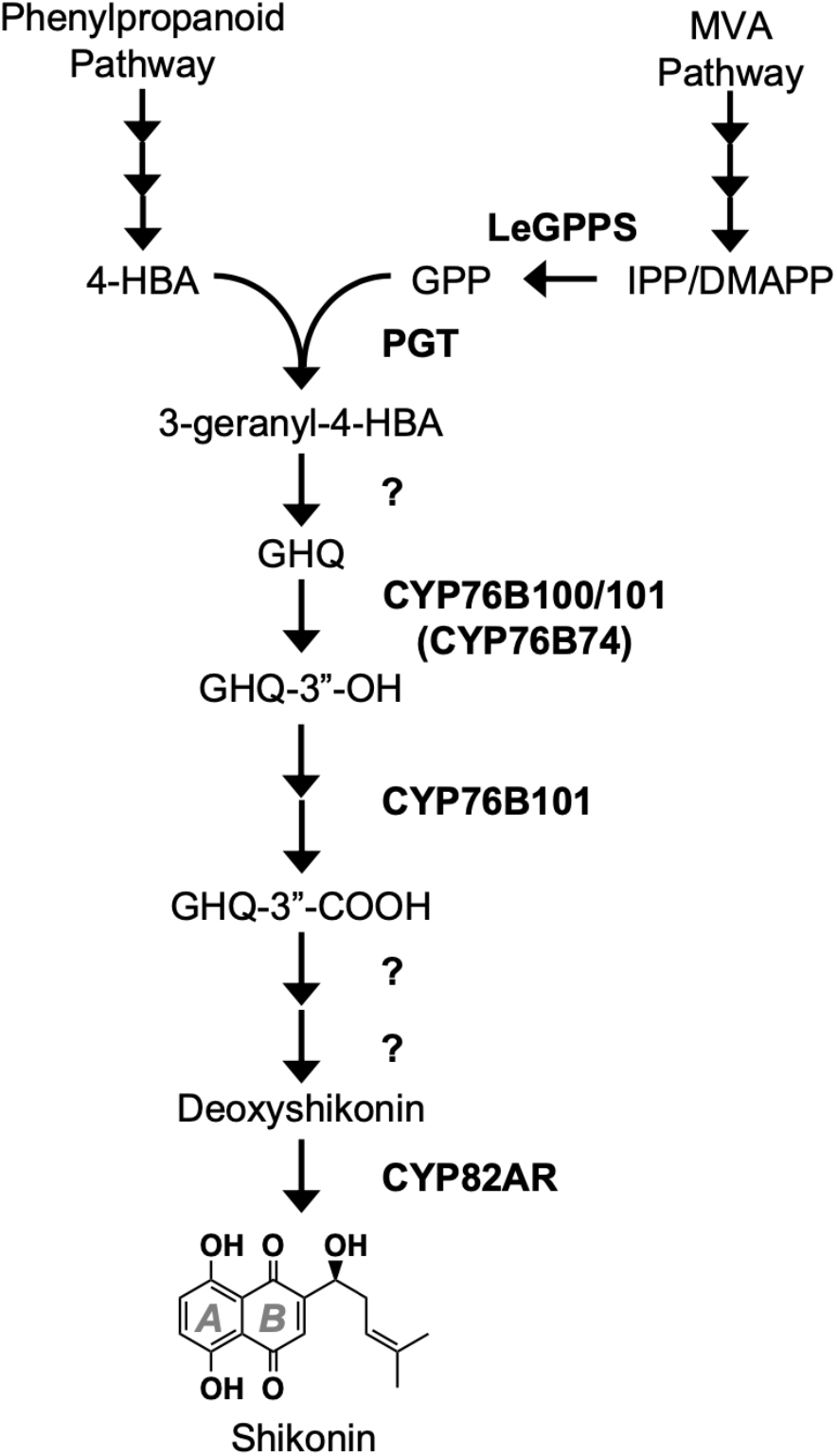
The shikonin metabolic network. Depicted is the current understanding of the enzymes and intermediates involved in synthesizing shikonin from precursors of the phenylpropanoid (4-HBA) and the MVA (GPP) pathways. Question marks indicate proposed steps lacking experimental evidence. Abbreviations: 4-HBA, 4-hydroxybenzoate; CYP76B74 (*Arnebia euchroma*) and CYP76B100/101 *(Lithospermum erythrorhizon)*, GHQ 3”-hydroxylase; CYP82AR, deoxyshikonin hydroxylase; DMAPP, dimethylallyl diphosphate; GHQ, geranylhydroquinone; GPP, geranyl diphosphate; GPPS, geranyl diphosphate synthase; IPP, isopentenyl diphosphate; MVA, mevalonic acid; PGT, p-hydroxybenzoate:geranyltransferase.

The structure of shikonin is comprised of a redox-active naphthazarin (5,6-dihydroxy-1,4-naphthoquinone) ring fused with a 1-hydroxy-4-methyl-3-pentenyl side chain (Fig. 1). The hydroxybenzene ring, ring *A*, of shikonin’s naphthazarin moiety is derived from L-phenylalanine via cinnamic acid and 4-hydroxybenzoate (4-HBA)^6, 7^. This is the same route predominantly responsible for forming the benzoquinone ring of ubiquinone (coenzyme Q) in plants^8, 9^. Many of the genes responsible for synthesis of the 4-HBA precursor of shikonin have already been cloned and investigated (e.g.^10–12^). In contrast, the genetic basis and regulation underlying the unique formation of the prenyl diphosphate precursor providing shikonin’s quinone ring, ring *B*, and its six-carbon atom isoprenoid side chain is not as well characterized.

The shikonin pathway begins with the conjugation of 4-hydroxybenzoate (4-HBA) and geranyl diphosphate (GPP) catalyzed by *<u>p</u>*-hydroxybenzoate:<u>g</u>eranyltransferase (PGT)^13^ to produce 3-geranyl-4-HBA^14^ (Fig. 1). GPP and other prenyl diphosphates are synthesized from the condensation of the five-carbon building blocks isopentenyl diphosphate (IPP) and dimethylallyl diphosphate (DMAPP). In plants, GPP is typically produced by plastidial GPP synthases (GPPSs) that catalyze the condensation of one IPP and one DMAPP derived from the methylerythritol phosphate (MEP) pathway localized in plastids^15^. Plants also produce IPP and DMAPP via the mevalonic acid (MVA) pathway, a route separated from the MEP pathway that is compartmentalized across the cytoplasm, endoplasmic reticulum, and peroxisomes^15–17^. The MVA pathway is generally considered to generate isoprenoid precursors for farnesyl diphosphate (FPP) synthases (FPPSs), which catalyze the condensation of one DMAPP with two IPP molecules to produce FPP and two molecules of pyrophosphate in the cytoplasm. Experimental evidence (reviewed in Widhalm and Rhodes^4^) long suggested that shikonin biosynthesis unconventionally relies on GPP produced by a cytoplasmic GPPS. The recent discovery and biochemical characterization of *L. erythrorhizon* GPPS (LeGPPS, Fig. 1)^18^ revealed that it is a neofunctionalized cytoplasmic farnesyl diphosphate synthase (FPPS) and that mutation(s) adjacent to the first aspartate-rich motif resulted in acquisition of GPPS activity^18^.

Previous analysis by our group of the *L. erythrorhizon* genome uncovered an evolutionary link between *PGT*s and the ubiquinone prenyltransferase gene, demonstrating that retrotransposition-derived gene duplication and subsequent neofunctionalization contributed to the evolution of *PGT* genes^19^. Coupled with the evolution of LeGPPS from a cytoplasmic FPPS^18^ and whole genome duplication (WGD) in the Boraginaceae^19, 20^, the evolutionary history of the shikonin pathway appears to be marked by several events of metabolic innovation. Taken together, this raises the prospect of additional evolutionary links between the shikonin and ubiquinone pathways and opens new questions about the metabolic intersection of the isoprenoid, phenylpropanoid, ubiquinone, and shikonin pathways in the Boraginaceae.

In this study, we investigated the metabolic connections linking shikonin biosynthesis with its precursor pathways by downregulating expression of *LeGPPS* and testing the capacity of the MEP and MVA pathways to supply GPP for shikonin production. We also explored whether network analysis of transcript abundances could identify genes coexpressed with *LeGPPS* and other established shikonin pathway genes. Integrative computational analyses of candidate genes identified by the model suggest likely metabolic roles for these genes and give insight into the evolution of metabolic innovation in the shikonin pathway. Our study provides evidence of crosstalk between the MVA, MEP, and phenylpropanoid pathways and reveals additional evolutionary links between shikonin and ubiquinone biosynthesis. Given the other links between specialized and primary quinone metabolism^21, 22^, the mechanistic insights uncovered here are expected to broadly guide investigation into the convergent evolution of specialized 1,4-naphthoquinone metabolism in plants.

## Results

### Cytoplasmic LeGPPS supplies GPP to the shikonin pathway using MVA pathway-derived IPP/DMAPP

To investigate the in vivo role of LeGPPS, which sits at the interface between the MVA, phenylpropanoid, and shikonin pathways, we knocked down expression of its encoding gene in *L. erythrorhizon* hairy roots. Several independent *LeGPPS*-RNAi (*LeGPPSi*) lines were generated, excised, and transferred to B5 selection media plates and subsequently screened for levels of total shikonins excreted into the growth media 4 d after transfer to M9 and darkness. Analysis of 10 independent *LeGPPS*-RNAi lines revealed that total shikonins were reduced by more than 95% compared to the lowest producing empty-vector control line (Fig. 2a). Further analysis of two independent lines, *LeGPPSi*-45 and *LeGPPSi*-75, revealed that *LeGPPS* expression was reduced by more than 95% compared to empty-vector control *EV-*26 without any affect on expression of the canonical plastidial GPPS gene *LepGPPS* (Fig. 2b). Re-analysis of total shikonins excreted from *LeGPPSi* lines 45 and 75 confirmed nearly 95% reduction compared to *EV*-26 (Fig. 2c), thus indicating that cytoplasmic LeGPPS is predominantly responsible for supplying GPP precursor to the shikonin pathway.

**Fig. 2.**
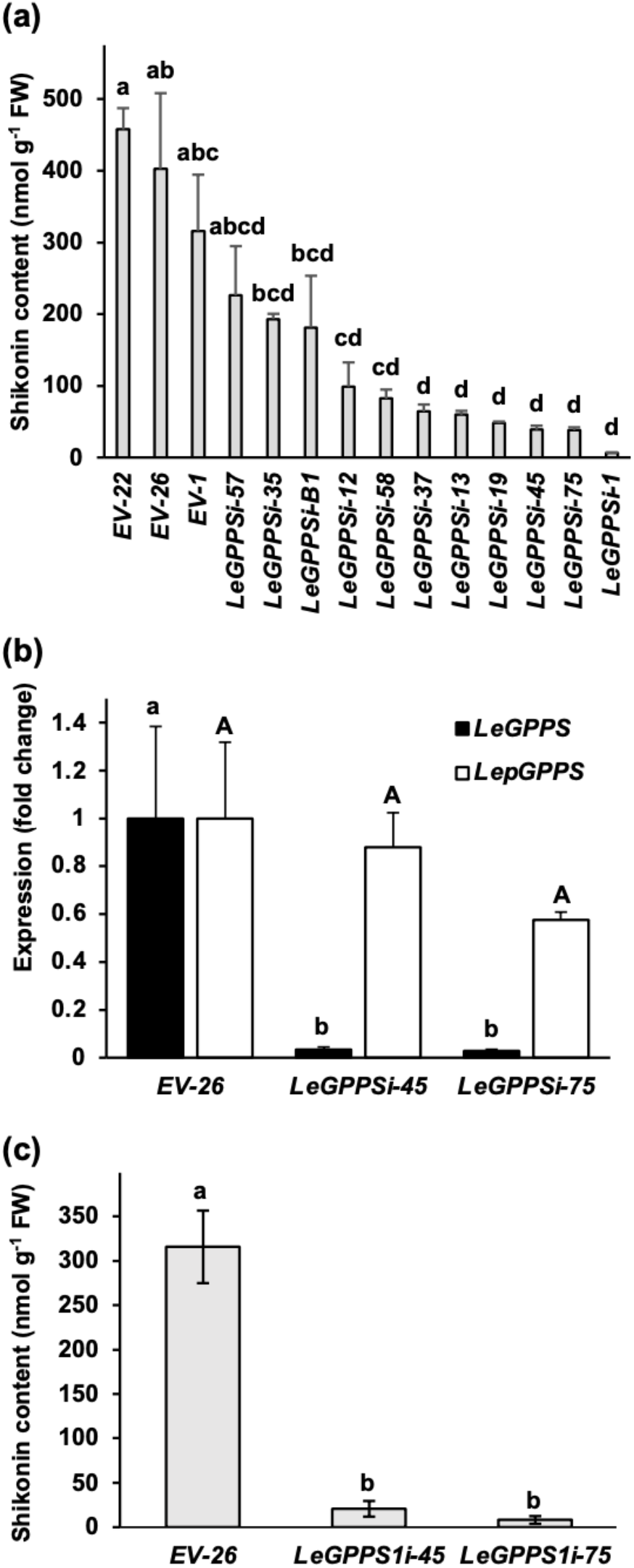
In vivo characterization of LeGPPS. Screening of *LeGPPS*-RNAi (*LeGPPS*i) lines based on total shikonin levels present in liquid culture media 3 d after transfer of 14-d-old hairy roots to M9 and darkness (a). Expression levels of *LeGPPS* and the canonical plastid-localized GPPS gene (*LepGPPS*) in hairy roots of two independent *LeGPPS*i lines compared to an empty-vector control line (*EV*-26) (b). Analysis of total shikonin in the same lines used to measure expression in panel b (c). All data are means ± SEM (n = 3-4 biological replicates). Different letters indicate significant differences via analysis of variance (ANOVA) followed by post-hoc Tukey test (α= 0.05). In panel b, lowercase and capital letters correspond to statistical comparisons for *LeGPPS* and *LepGPPS* expression, respectively.

Concievably, MEP pathway-derived GPP could contribute to the shikonin pathway if it or MEP pathway-derived IPP/DMAPP were exported from the plastid to the cytoplasm and used as substrate by LeGPPS. To test for MEP pathway involvement in shikonin production we carried out two inhibitor experiments on the *EV*-26 and *LeGPPSi*-45 lines (Fig. 1). We predicted that if the MVA pathway is predominantly responsible for supplying IPP/DMAPP to LeGPPS, then treatment with the MVA pathway inhibitor mevinolin should decrease shikonin accumulation in *EV*-26 lines but not in the *LeGPPSi*-45 RNAi line. Indeed, total shikonins produced by mevinolin-treated *EV*-26 lines were reduced by 76% compared to those in the *EV*-26 control lines (Fig. 3a), while shikonins in mevinolin-treated *LeGPPSi*-45 lines were unchanged compared to the *LeGPPSi*-45 control lines (Fig. 3b). If the MEP pathway does not supply IPP/DMAPP precursor to the shikonin pathway, we expected no change in shikonin accumulation in *EV*-26 lines treated with the MEP pathway inhibitor fosmidomycin compared to controls. If, however, the MEP pathway is contributing to the remaining shikonin produced by *LeGPPSi*-45 lines, treatment with fosmidomycin should further reduce shikonin accumulation compared to *LeGPPSi*-45 controls. Instead, we observed that shikonin production increased by 73% and 108%, respectively, in *EV*-26 and *LeGPPSi*-45 lines treated with fosmidomycin compared to their corresponding controls (Fig. 3a,b). This result points to crosstalk between the MEP and MVA pathways such that when flux through the MEP pathway is impaired, flux through the MVA pathway is increased. Taken together, our genetic and inhibitor studies support the work of Ueoka *et al.*^18^ by showing that LeGPPS is required for shikonin formation and that the MEP pathway does not supply IPP or DMAPP substrates, or direct GPP precursor to the shikonin pathway.

**Fig. 3.**
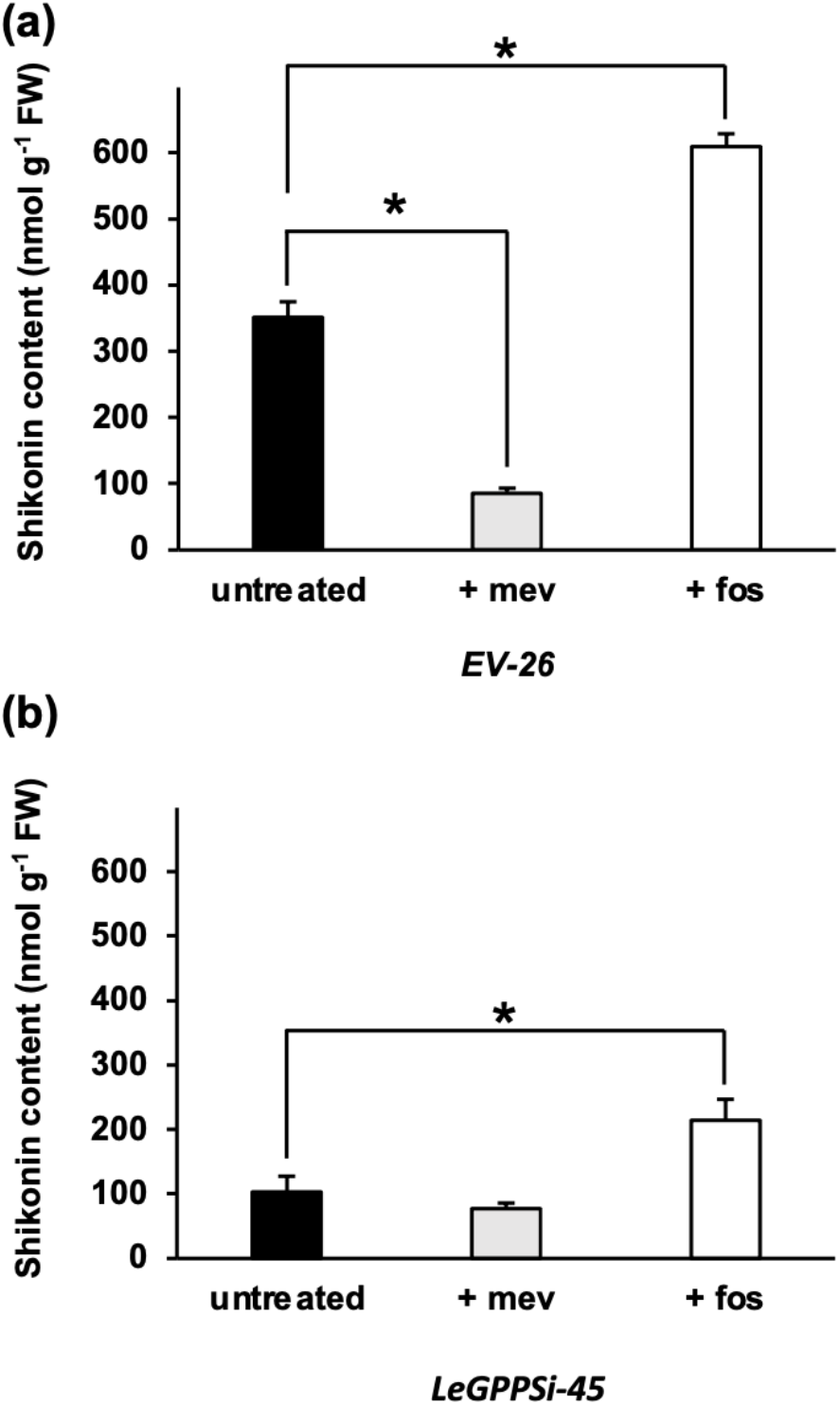
Effect of MVA and MEP pathway-specific inhibitors on formation of total shikonins. Total shikonin levels present in liquid culture media were measured in empty-vector control line 26 (*EV*-26) (a) and *LeGPPS* RNAi line 45 (*LeGPPSi*-45) (b) following mock treatment or treatment with the MVA pathway inhibitor mevinolin (+ mev) or the MEP pathway inhibitor fosmidomycin (+ fos). Inhibitor treatments were administered immediately upon transfer of 14-d-old hairy roots to M9 and darkness. Total shikonins were measure at 6 d after transfer of 14-d-old hairy roots to M9 and darkness. All data are means ± SEM (n = 3–4 biological replicates). Statistically significant differences are indicated (*P < 0.05, Student’s *t* test).

### Downregulation of *LeGPPS* reveals crosstalk between phenylpropanoid and isoprenoid metabolism

The observed increase in shikonin content in *LeGPPSi*-45 RNAi lines treated with the MEP pathway inhibitor fosmidomycin (Fig. 3b) led us to hypothesize that the smaller pool size of shikonin in *LeGPPS*-RNAi lines (Fig. 2) may be due, in part, to an upstream effect on the MVA pathway. To investigate if MVA pathway gene expression is changed, we performed RNA-seq analysis of *LeGPPSi*-45 lines compared to *EV-26* control. Our analysis showed 6,115 differentially expressed genes (DEGs); 2,903 genes were significantly overexpressed in *LeGPPSi*-45 lines compared to *EV*-26 while 3,212 were significantly underexpressed (Table S1), including shikonin pathway genes *LePGT1*, *LePGT2*, *CYP76B100*, and *CYP82AR* (Fig. S1). Kyoto Encyclopedia of Genes and Genomes (KEGG) term enrichment analysis of genes underexpressed in the *LeGPPSi*-45 line revealed an enrichment of genes involved in various metabolic pathways connected to shikonin metabolism (BH-adjusted *p*-value < 0.05; Fig. 4a, Fig. S2). The category “monoterpenoid biosynthesis,” which encompases metabolic genes downstream of GPP was significantly enriched among underexpressed genes. The KEGG category “terpenoid backbone biosynthesis,” which contains the MVA and MEP pathway genes, was not significantly enriched (BH-adjusted *p*-value = 0.097; Fig. S2). Yet, 11 of the 17 MVA pathway genes involved in IPP biosynthesis were found to be significantly underexpressed in *LeGPPSi*-45. This included six of the eight genes encoding 3-hydroxy-3-methylglutaryl-CoA reductase (HMGR), which is generally considered to catalyze the rate-limiting step of the MVA pathway (Fig. 4a, Table S1)^23^. This suggests that lower expression of upstream MVA pathway genes may have contributed to reduced shikonin production in *LeGPPS*-RNAi lines (Fig. 2). These data also point to an unknown factor connecting downregulation of *LeGPPS* with reduced expression of upstream MVA pathway genes.

**Fig. 4.**
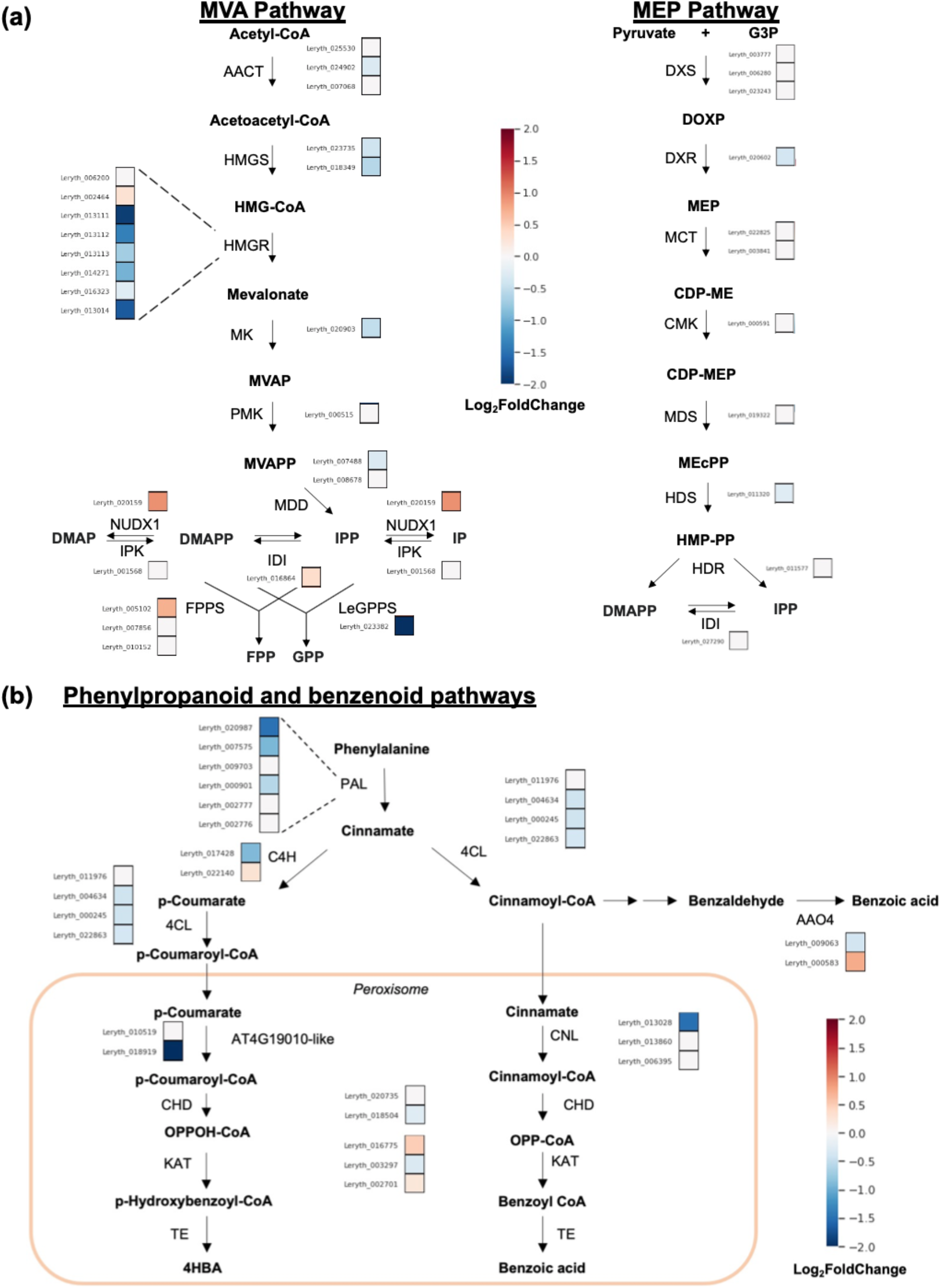
Effect of *LeGPPS* RNAi downregulation on expression of MVA, MEP, phenylpropanoid, and benzenoid pathway genes. The average log_2_fold-change in expression for each gene in *LeGPPSi*-45 lines compared to *EV-26* lines in the mevalonic acid (MVA) and methylerythritol phosphate (MEP) pathways (a) and in the phenylpropanoid and benzenoid pathways (b) are shown. Abbreviations: 4CL, 4-coumarate CoA-ligase; AACT, acetoacetyl-CoA thiolase; AA04, Arabidopsis Aldehyde Oxidase 4; C4H, cinnamate 4-hydroxylase; CDP-ME, 4-Diphosphocytidyl-2-C-methylerythritol; CDP-MEP, 4-Diphosphocytidyl-2-C-methylerythritol 2-phosphate; CHD, cinnamoyl-CoA hydratase/dehydrogenase; CMK, 4-(cytidine 5ʹ -diphospho)-2-C-methyl-D-erythritol kinase; CoA, coenzyme A; DMAP, dimethylallyl phosphate; DMAPP, dimethylallyl diphosphate; DOXP, 1-deoxy-D-xylulose 5-phosphate; DXR, 1-deoxy-D-xylulose 5-phosphate reductoisomerase; DXS, 1-deoxy-D-xylulose 5-phosphate synthase; FPP, farnesyl diphosphate; FPPS, farnesyl diphosphate synthase; G3P, D-glyceraldehyde 3-phosphate; GPP, geranyl diphosphate; GPPS, geranyl diphosphate synthase; HDR, (E)-4-hydroxy-3-methylbut-2-enyl diphosphate reductase; HDS, (E)-4-hydroxy-3-methylbut-2-enyl diphosphate synthase; HMG-CoA, 3-hydroxy-3-methylglutaryl-CoA; HMGR, 3-hydroxy-3-methylglutaryl-CoA reductase; HMGS, 3-hydroxy-3-methylglutaryl-CoA synthase; HMP-PP, (E)-1-hydroxy-2-methylbut-2-enyl 4-diphosphate; IDI, isopentenyl diphosphate isomerase; IP, isopentenyl phosphate; IPK, isopentyl phosphate kinase; IPP, isopentenyl diphosphate; KAT, 3-ketoacylthiolase 1; MCT, 2-C-methyl-D-erythritol 4-phosphate cytidylyltransferase; MDD, mevalonate diphosphate decarboxylase; MDS, 2-C-methyl-D-erythritol 2,4-cyclodiphosphate synthase; MEcPP, methylerythritol cyclodiphosphate; MK, mevalonate kinase; MPD, phosphomevalonate decarboxylase; MVAP, mevalonate 5-phosphate; MVAPP, mevalonate diphosphate; NUDX1, Nudix enzyme 1; OPP-CoA, 3-oxo-3-phenylpropionoyl-CoA; PAL, L-phenylalanine ammonia lyase; PMK, phosphomevalonate kinase; PXA1, peroxisomal ABC transporter 1; TE, thioesterase.

The KEGG pathway analysis also revealed that genes involved in “phenylpropanoid biosynthesis” and “ubiquinone and other terpenoid-quinone biosynthesis” were enriched in those underexpressed in *LeGPPSi*-45 (Fig. S2). This is noteworthy because the phenylpropanoid pathway supplies *p*-coumaroyl-CoA to make the 4-HBA precursor that becomes the hydroxybenzene ring, ring *A*, of shikonin’s naphthazarin moiety (Fig. 1) and of ubiquinone’s benzenoid moiety^8^. Further examination of genes underexpressed in *LeGPPSi*-45 showed that several genes in the core phenylpropanoid pathway are underexpressed, including multiple genes encoding phenylalanine ammonia-lyases (PALs) (Fig. 4b, Table S1). Moreover, one copy of the At4g19010-like peroxisomal *p*-coumarate-CoA ligase genes (Leryth_018919) was significantly underexpressed (Fig. 4b, Table S1). In *Arabidopsis thaliana*, it was demonstrated that At4g19010 is responsible for activating the propyl side chain of *p*-coumarate for *β*-oxidative shortening to supply 4-HBA precursor for ubiquinone biosynthesis^8^. These results suggest that, like the MVA pathway, an unknown factor links downregulation of *LeGPPS* to reduced expression of phenylpropanoid and benzenoid pathway genes.

Decreased accumulation of transcripts encoding core phenylpropanoid and *β*-oxidative benzenoid biosynthetic genes (Fig. 4b) raises the possibility that 4-HBA availability might also limit shikonin production in *GPPSi*-RNAi lines (Fig. 2). To test this, shikonin levels were determined in *EV*-26 and *LeGPPSi*-45 lines supplied with exogenous 4-HBA. The amount of shikonin produced, however, remained unchanged compared to the unfed controls (Fig. S3) suggesting that 4-HBA availability does not limit shikonin production in *LeGPPS* knockdown lines. Taken together, the in vivo investigation of *LeGPPS* demonstrates that in addition to LeGPPS being involved in shikonin biosynthesis, the expression of *LeGPPS* is highly connected to other genes in the larger shikonin metabolic network including those in the shikonin, MVA, phenylpropanoid, and benzenoid pathways.

### Coexpression network analysis recovers known shikonin pathway gene associations and predicts new connections

We hypothesized that *LeGPPS* and other known shikonin biosynthesis genes would appear as hub genes that we could use to identify coexpressed genes with roles in shikonin biosynthesis. To construct a transcriptional network model with a high likelihood of recovering the shikonin biosynthetic pathway as a module, we used publicly available comparative RNA-seq experiments from tissues and conditions divergent in their shikonin levels (NCBI Sequence Read Archives PRJNA596998^19^ and PRJNA331015). These included whole *L. erythrorhizon* root tissue versus above ground tissue; root periderm versus root vascular (inner) tissue; and hairy root cultures grown in M9 media in the dark versus roots grown in B5 media in light conditions. In all three experiments, the former tissue or condition in each comparison was previously shown to contain higher *LePGT1* expression and shikonin content^19^. Like LeGPPS, LePGT1 functions at the interface of the phenylpropanoid, MVA, and shikonin pathways (Fig. 1). Therefore, we constructed the model based on the hypothesis that genes involved in the shikonin pathway and upstream metabolism would also be more highly expressed in the same tissues or conditions as *LePGT1*.

One potential source of noise in coexpression analyses is the inclusion of genes that are either not expressed or are constitutively expressed at a constant level across all conditions. These genes may appear significantly coexpressed with other genes in the dataset artifactually^24^. Although this is less of a concern when working with dozens or hundreds or RNA-seq samples^25^, our dataset consisted of only 14 RNA-seq samples across six total tissues or conditions. To control for this source of false positive coexpression, we only included genes likely to be DEGs in at least one of the three comparisons at a false discovery rate (FDR) cutoff of ≤ 0.1. A total of 8,680 transcripts were included using this approach (Table S2). Within this set of transcripts, 23.9% (2,077) were overexpressed in whole root; 37.9% (3,290) were overexpressed in root periderm; and 21.6% (1,876) were overexpressed in hairy roots sampled in the dark (Table S3). The overlap of all three comparisons contained 374 genes that were overexpressed in the shikonin accumulating condition (Fig. 5a). These included several genes already implicated in shikonin biosynthesis: *LePGT1*, *LePGT2*, two additional *PGT*-like genes^19^, *LeGPPS*^18^*, CYP82AR2*^26^*, L. erythrorhizon pigment callus-specific gene 2* (*LePS-2*)^27^, and *LeMYB1*^28^ (Table S4).

**Fig. 5.**
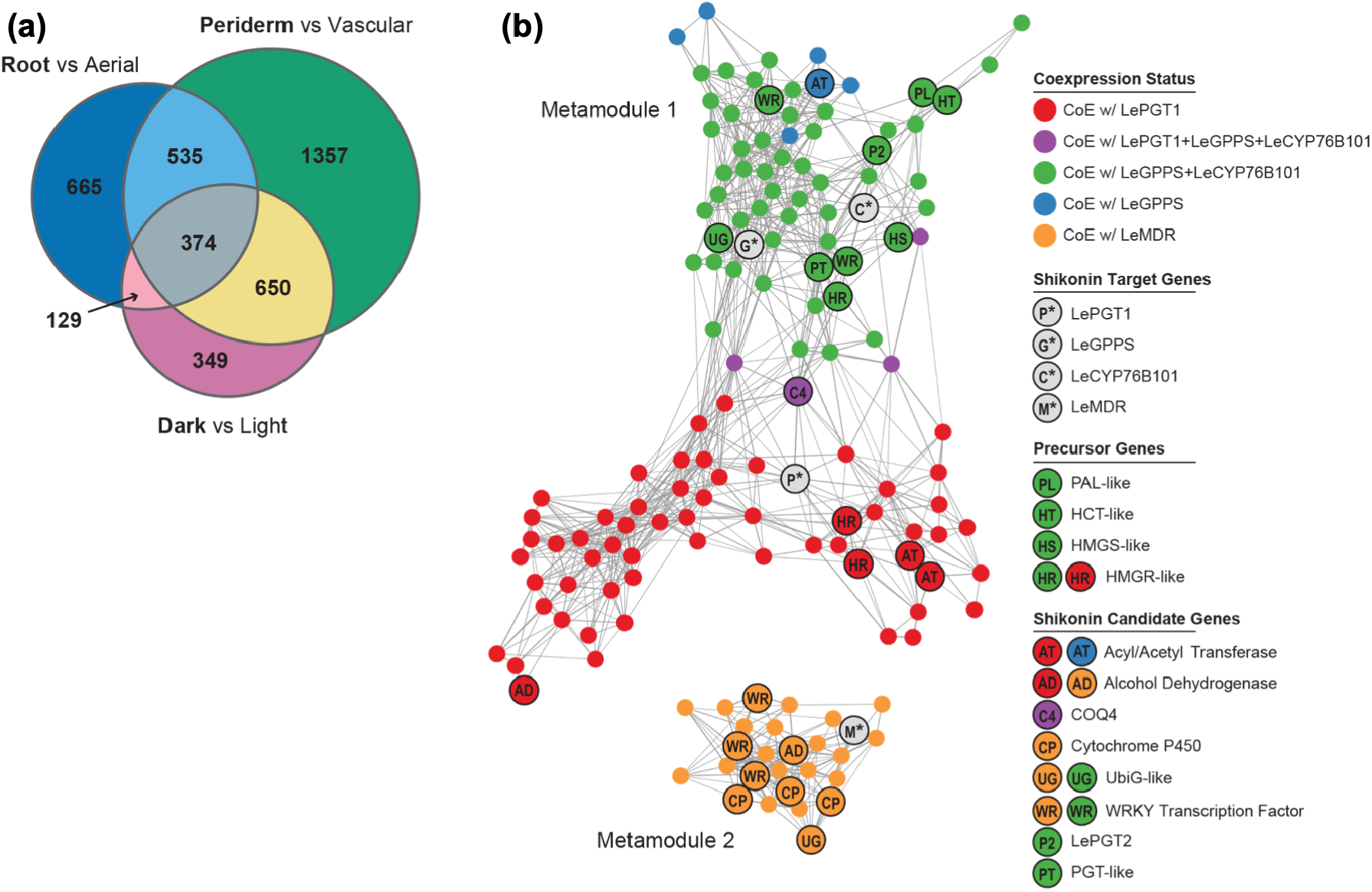
Analysis of gene expression in *L. erythrorhizon*. Venn diagram showing the overlap of genes that are significantly overexpressed in conditions where shikonin is abundant (bold) (a). Network map of genes coexpressed with target genes *LePGT*, *LeGPPS*, *LeCYP76B101*, and *LeMDR* using the N2 global coexpression network (b). Nodes in the map represent genes, and edges connecting two genes represent the weight (transformed MR score) for the association. Genes are colored according to its coexpression status with known shikonin genes (grey). Network maps were drawn using a Fruchterman-Reingold force-directed layout using the edge-weighted spring embedded layout in cytoscape (https://cytoscape.org).

The 8,680 DEGs were used as input for a global coexpression network analysis (Table S5). Pairwise measurements of gene coexpression were specified as mutual ranks (MRs), which are calculated as the geometric mean of the rank of the Pearson’s correlation coefficient (PCC) of gene A to gene B and the PCC rank of gene B to gene A^29^. Ranking the PCCs in this manner has been shown to improve the recovery of known pathways as discrete subgraphs in global coexpression networks^30^. We constructed four MR-based networks (N1-N4), using different coexpression thresholds for assigning edge weights (i.e., connections) between nodes (i.e., genes) in the network. Networks were ordered by size (i.e., total number of edges between nodes), such that N1 represents the smallest network and N4 represents the largest network. Graph-clustering implemented by ClusterONE^31^ was used to discover coexpressed subgraphs (hereafter referred to as gene modules) within the global networks (Dataset S1). The benefit of using ClusterONE over other graph-clustering methods, e.g. MCL^32^ is its capacity to assign genes to multiple overlapping modules, which is more reflective of complex biological networks. We chose to focus our analysis on four target genes based on evidence of their involvement in shikonin metabolism: *LeGPPS*^18^ (Fig. 2), *LePGT1*^19^, *LeCYP76B101*^33, 34^, and *LeMDR*^35^. Because ClusterONE modules can overlap, each target gene was assigned to multiple modules within the larger networks. For example, *LePGT1* was found in 3, 2, 3, and 6 different modules in network N1, N2, N3, and N4, respectively (Dataset S1). To address this redundancy, we collapsed all modules within a network that contained one or more of the four target shikonin metabolic genes into non-intersecting metamodules (Fig. 5b; Fig. S4)^25^. Collectively, these metamodules are models, which we refer to as shikonin metabolic subnetworks.

The number of genes recovered in the shikonin metabolic subnetworks varied from 102, 152, 359, and 1,268 genes in networks N1, N2, N3, and N4, respectively (Tables S6-S9). We focused our subsequent analyses on the N2 network, which contained a large number of candidate genes to investigate while also limiting the number of peripheral genes that appeared only weakly connected to shikonin biosynthesis (Fig. S4). The N2 shikonin metabolic subnetwork was comprised of two metamodules (Fig. 5b). The first N2 metamodule contained 125 genes including *LeGPPS*, *LePGT1*, and *LeCYP76B101*; whereas, the second metamodule contained 27 genes including *LeMDR*. To be considered coexpressed in our analysis two genes must have at least one shared module within the larger metamodule. For example, *LeGPPS* and *CYP76B101* were coexpressed with one another, being members of three shared modules: N2M94, N2M298, and N2M317 (Dataset S1). Within metamodule 1, 60 genes were coexpressed with *LeGPPS* and *CYP76B101*; 6 genes were uniquely coexpressed with *LeGPPS*; and 59 genes were uniquely coexpressed with *LePGT1* (Fig. 5b; Table S7). Four genes (Leryth_014746, Leryth_025160, Leyrth_004583, Leryth_002195) coexpressed with all three *LeGPPS*, *LePGT1*, and *LeCYP76B101* (Fig. 5b; Table S7). Although the genes of metamodule 2, including *LeMDR*, were not coexpressed with the three other target shikonin genes in network N2, the two larger networks N3 and N4 did show a small amount of overlap (Fig. S4).

In agreement with previous studies^36, 37^, six genes were recovered in N2 metamodule 1 encoding enzymes with annotations related to the phenylpropanoid and MVA pathways including PAL, HCT, HMGS, and HMGR (Fig. 5b; Table S7). Past studies have identified additional candidate genes possibly involved in the shikonin pathway including *LePS-2*^27^, *LeACS-1*^38^, *LeMYB1*^28^, and *LeDI-2*^39^. Of these, only *LePS-2* was coexpressed with any validated shikonin biosynthetic genes, being coexpressed with *LePGT1*, *LeGPPS*, and *LeCYP76B101* in the larger N3 and N4 networks (Tables S8,S9).

The N2 shikonin subnetwork was enriched in 62 Gene Ontology (GO) categories (BH-adjusted *p*-value < 0.05; Table S10) including broad enzymatic categories such as GO:0016491 oxidoreductase activity (18 genes), and GO:0016740 transferase activity (36 genes). Another enriched category, ATPase-coupled intramembrane lipid transport activity (2 genes; Leryth_023505, Leryth_019206) is of high interest because the previous implication of an ARF/GEF-like system required for shikonin transport^40^.

To identify shared 5’ cis regulatory regions among the coexpressed genes, we performed a motif enrichment analysis on the genes of the N2 shikonin subnetwork using Motif Indexer^41^. The most overrepresented motif within the upstream region of shikonin subnetwork genes was AmrGTCwA (*p*-value = 9.67×10^-10^; FDR = 0.007; Table S11), the reverse compliment of which (TwGACykT) is similar to the canonical W-box element sequence motif (T)TGAC(C/T) recognized by the WRKY family of transcription factors^42^. Of the 152 genes in the N2 shikonin subnetwork, 47.37% of genes (N = 72) contained this motif including all four target genes: *LePGT1, LeGPPS*, *LeMDR*, and *CYP76B101*. Five WRKY transcription factors were identified in the N2 shikonin subnetwork (Fig. 5b) two of which (Leryth_027519 and Leryth_002564) were also significantly overexpressed in all three conditions where shikonin was abundant (Table S7).

### The evolutionary history of *LeGPPS* suggests an expanded role of the FPPS gene family in specialized metabolism

Previous work demonstrated that *LeGPPS* encodes an enzyme having GPPS-like activity but is a member of the *FPPS* gene family^18^. To better understand the evolutionary history of *LeGPPS*, we reconstructed the phylogeny of the *FPPS* gene family using homologous sequence groups downloaded from the PLAZA 4.0 database (Table S12)^43^. In addition to *L. erythrorhizon*, we included in our analysis de novo transcriptome-based proteomes from 18 additional Boraginales species including three other shikonin producing plants (*Echium plantagineum*, *Arnebia euchroma*, and *Lithospermum officinale*), one additional Boraginaceae that does not produce shikonin (*Mertensia paniculata*), and 14 additional Boraginales species that do not produce shikonin (Table S13)^19, 44^. The Boraginales contain two distinct subfamilies in the *FPPS* gene family phylogeny (Fig. 6). Subfamily I contains *LeGPPS* and was present in 17 of the 19 Boraginales in the analysis, including all four shikonin-producing species (Fig. 6). Subfamily I was absent in *Heliotropium karwinsky* and *Heliotropium* sp. Subfamily II, the canonical FPPS group, contains *LeFPPS1* (Leryth_005102)and 29 other sequences. Subfamily II was present in all four shikonin-producing species and absent in *Heliotropium calcicole* and *Heliotropium texanum*. The absence of subfamily I or II in some *Heliotropium* species is likely artifactual due to these gene sets being transcriptome derived. Subfamily II also contained two additional genes from *L. erythrorhizon*, Leryth_007856 (referred to as *LeFPPS2* by Ueoka *et al.*^18^) and Leryth_010152 (hereafter *LeFPPS3*) (Fig. 6).

**Fig. 6.**
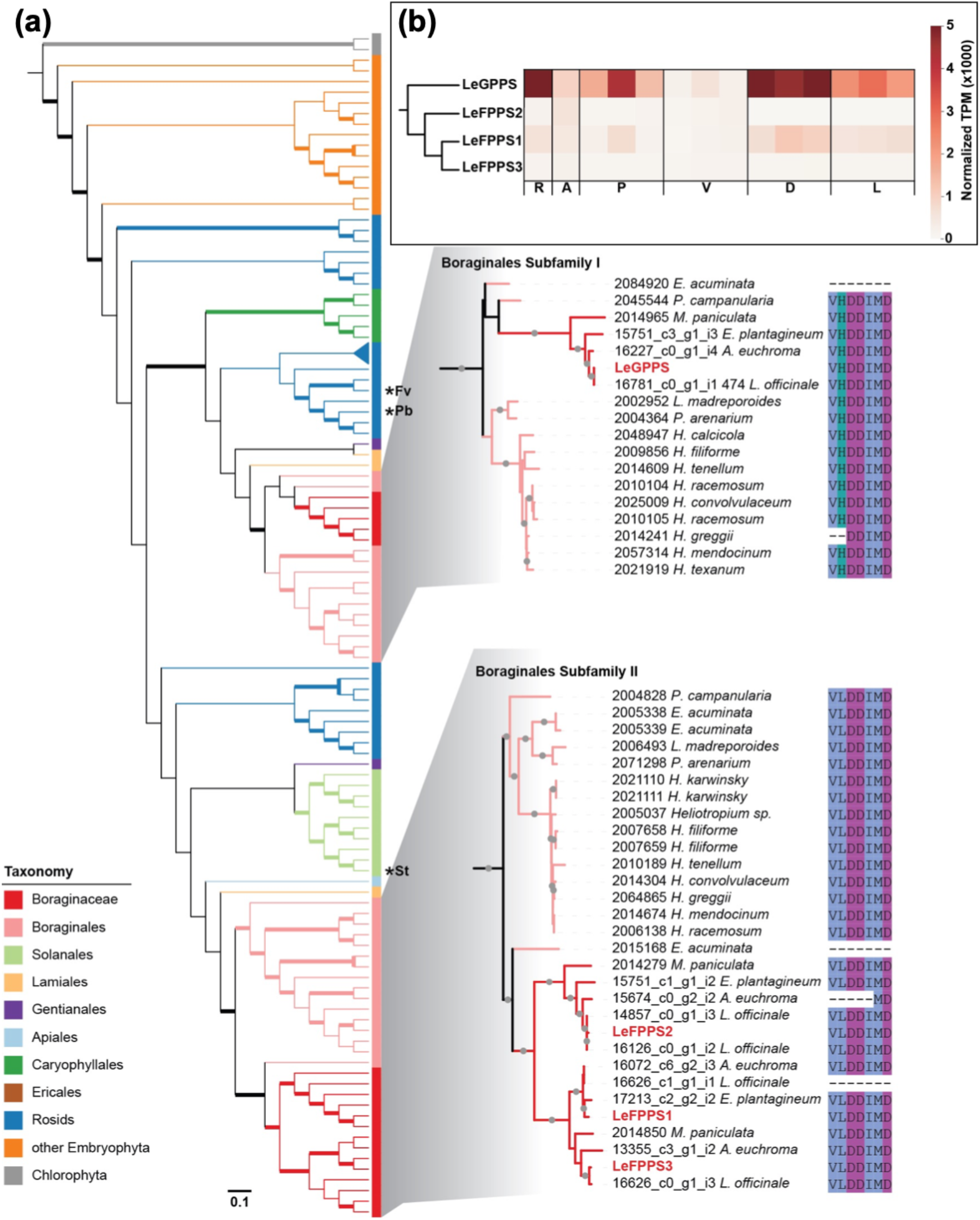
Phylogenetic analysis and expression profile of FPPS genes. Maximum likelihood tree of FPPS gene family in green plants (a). In the full cladogram, left, nodes with IQ-TREE support values > 95 are bolded. The branches and outer color bar are color-coded to match the taxonomic classification of each sequence. The tree is rooted on Chlorophytes. Non-Boraginales sequences containing a Histidine residue adjacent to the conserved Asp-rich motif are indicated by an asterisk (Fv, *Fragaria vesca*; Pb, *Pyrus bretschneideri*; St, *Solanum tuberosum*). Detailed phylograms of the two Boraginales clades, right, along with an alignment of Asp-rich motif (residues 309-315). Sequences from *L. erythrorhizon* are red. Nodes in phylograms with support values > 95 are indicated by the grey circles. The full phylogram for the entire FPPS family with branch lengths and leaf labels is available in the Supplemental. Heatmap showing the gene expression pattern of *L. erythrorhizon* FPPS genes in whole roots (R), arial tissue (A), root periderm (P), root vascular (V), hairy root grown in dark (D), and hairy root grown in the light (L) (b). The cladogram (left) shows evolutionary relationship between FPPS genes according to the overall maximum likelihood phylogeny in part (a).

We searched for shared synteny between genome assembly contigs containing *FPPS* genes in *L. erythrorhizon* to investigate whether whole genome duplication (WGD) was involved in the evolution of the subfamily I. A WGD is proposed for the Boraginaceae roughtly 25 MYA^20^ and *L. erythrorhizon* and *E. plantagineum* have similar distributions of synonymous substitution (Ks) between syntenic paralogs at 0.45 and 0.417, respectively^19, 20^. The contigs containing *LeFPPS1* and *LeFPPS3* (Fig. S5) were syntenic and the syntelogs in these two contigs have a median Ks value of 0.484 (Table S14). The median Ks of this syntenic block is similar to the peaks in Ks distribution described by Auber *et al.*^19^ and Tang *et al.*^20^, consistent with the Boraginaceae WGD giving rise to *LeFPPS1* and *LeFPPS3*. In contrast, the lack of shared synteny between *LeGPPS* and any of the three genes in the FPPS group suggests that *LeGPPS* did not arise via WGD. Intron position is conserved between *LeGPPS* and the three *FPPS* genes, with only *LeFPPS3* showing some divergent intron positioning toward its 3’ end (Fig. S6), which is consistent with segmental duplication or DNA transposition giving rise to the *LeGPPS* homolog rather than retrotransposition.

Previous work by Ueoka *et al.*^18^ demonstrated that the histidine (His) residue adjacent to the first aspartate-rich motif in LeGPPS was responsible for its GPPS-like activity. Examination of our *FPPS* gene family sequence alignment shows that this His residue is present in all sequences of the GPPS group (subfamily I), with the exception of two transcriptome-derived sequences from two non shikonin-producing species *Ehretia acuminata* and *Heliotropium greggii* that are both missing this region (Fig. 6). In contrast, all sequences in the FPPS group (subfamily II) contain the canonical leucine (Leu) residue adjacent to the Asp-rich motif, with the exception of three transcriptome-derived sequences that are missing the region (Fig. 6). We identified a His residue in place of Leu in three additional sequences from *Fragaria vesca* (woodland strawberry), *Pyrus bretschneideri* (Chinese white pear), and *Solanum tuberosum* (potato) (Fig. 6; Fig. S7). Like the Boraginales, each of these three species maintained a second FPPS gene that retains the canonical Leu adjacent to the Asp-rich motif (Fig. S7). A similar observation was made with FPPS homologs from *Fragaria x ananassa* (strawberry), *Malus domestica* (apple), and *Prunus persica* (peach)^18^. Thus, the recruitment of a cytoplasmic FPPS to function as a GPPS convergently evolved multiple times in plants and has likely contributed to the diversification of plant terpenoid metabolism.

### Shikonin pathway gene candidates provide insights into specialized metabolic innovation in the Boraginaceae

We extended our phylogenetic analysis to additional shikonin gene candidates (Table 1). We first considered gene candidates that could be responsible for missing enzymes in the shikonin pathway. It is estimated that 97% of cytochromes P450 in plants are associated with specialized metabolic pathways^45^. Therefore, considering that missing steps in the shikonin pathway require decarboxylation, hydroxylations, or carbon-carbon ring closure, we examined cytochromes P450 in the coexpression network. In addition to *LeCYP76B101*, which was used as a known target in metamodule construction, three additional cytochromes P450 were recovered in the N2 shikonin subnetwork, all three of which were coexpressed with *LeMDR* metamodule 2 (Fig. 5b; Table S7). None of the three additional cytochromes P450 correspond to the *LeCYP82AR2* recently described to catalyze deoxyshikonin hydroxylation in vitro^26^. Although *LeCYP82AR2* (Leryth_026973, Table S3) was not recovered as a candidate in the N1 or N2 shikonin subnetworks, it was coexpressed with *LePGT1*, *LeGPPS*, and *LeCYP76B101* in our N3 network (Fig. S4c) and was also overexpressed in all shikonin-abundant conditions (Table S3).

**Table 1.** Shikonin pathway gene candidates identified via coexpression network analysis

| Gene | InterPro | Network | Coexpressed with | DE DvL | DE PvV | DE RvL |
| --- | --- | --- | --- | --- | --- | --- |
| Leryth_021809 | Cytochrome P450 | N2 | *LeMDR* | yes | yes | yes |
| Leryth_001242 | Cytochrome P450 | N2 | *LeMDR* |  | yes | yes |
| Leryth_000257 | Cytochrome P450 | N2 | *LeMDR* |  | yes |  |
| Leryth_002195 | Ubiquinone biosynthesis protein COQ4 | N2 | *LeGPPS*, *LePGT*, *CYP76B101* | yes | yes | yes |
| Leryth_019821 | Ubiquinone biosynthesis O-methyltransferase COQ3 | N2 | *LeMDR* |  | yes | yes |
| Leryth_021171 | Ubiquinone biosynthesis O-methyltransferase COQ3 | N2 | *LeGPPS*, *CYP76B101* | yes | yes |  |
| Leryth_001358 | Prephenate dehydrogenase | N2 | *LePGT* | yes | yes |  |
| Leryth_020454 | Quinoprotein glucose/sorbose dehydrogenase | N4 | *LeGPPS*, *LePGT*, *CYP76B101* | yes | yes | yes |
| Leryth_012925 | Chloramphenicol acetyltransferase-like | N2 | *LePGT* |  | yes | yes |
| Leryth_015823 | Chloramphenicol acetyltransferase-like | N2 | *LeGPPS*, *CYP76B101* |  | yes | yes |

**Table 1** 1114 Shikonin pathway gene candidates identified via coexpression network analysis
| Gene | InterPro | Network | Coexpressed with | DE DvL | DE PvV | DE RvL |
| --- | --- | --- | --- | --- | --- | --- |
| Leryth_021809 | Cytochrome P450 | N2 | *LeMDR* | yes | yes | yes |
| Leryth_001242 | Cytochrome P450 | N2 | *LeMDR* |  | yes | yes |
| Leryth_000257 | Cytochrome P450 | N2 | *LeMDR* |  | yes |  |
| Leryth_002195 | Ubiquinone biosynthesis protein COQ4 | N2 | *LeGPPS, LePGT, CYP76B101* | yes | yes | yes |
| Leryth_019821 | Ubiquinone biosynthesis O-methyltransferase COQ3 | N2 | *LeMDR* |  | yes | yes |
| Leryth_021171 | Ubiquinone biosynthesis O-methyltransferase COQ3 | N2 | *LeGPPS, CYP76B101* | yes | yes |  |
| Leryth_001358 | Prephenate dehydrogenase | N2 | *LePGT* | yes | yes |  |
| Leryth_020454 | Quinoprotein glucose/sorbose dehydrogenase | N4 | *LeGPPS, LePGT, CYP76B101* | yes | yes | yes |
| Leryth_012925 | Chloramphenicol acetyltransferase-like | N2 | *LePGT* |  | yes | yes |
| Leryth_015823 | Chloramphenicol acetyltransferase-like | N2 | *LeGPPS, CYP76B101* |  | yes | yes |

One of the three cytochromes P450 identified was Leryth_021809, which encodes a CYP76B6-like enzyme and was significantly overexpressed in all shikonin-abundant conditions (Table S3). Other *CYP76B* genes, including *CYP76B74* in *A. euchroma* ^34^ and *CYP76B100/101*^33^ in *L. erythrorhizon* (Fig. 1), have already been implicated in oxidative reactions in shikonin biosynthesis, but the evolutionary relationships between these genes has been unclear. A phylogeny of *CYP76B6*-like genes reveals that *AeCYP76B74* and *LeCYP76B101* are orthologs (Fig. S8). *LeCYP76B100* is the mostly closely related paralog to *LeCYP76B101* but groups more closely to other sequences in *A. euchroma, E. plantagineum*, and *M. paniculata* (Fig. S8). This indicates that the gene duplication event that gave rise to *LeCYP76B100/101* occurred in the last common ancestor of these Boraginaceae species. Leryth_021809, the additional CYP76B6-like gene recovered in the coexpression analysis, is within a separate clade that has expanded within shikonin producing species. The closest homolog in *M. paniculata* (the only Boraginaceae in our analysis that does not produce shikonin) groups closer to a different cytochrome P450 in *L. erythrorhizon* (Leryth_021691; Fig. S8).

A phylogeny of sequences homologous to the second cytochrome P450 candidate (Leryth_001242), which encodes a CYP76A2-like enzyme, also shows an expansion of gene copies in the Boraginaceae (Fig. S9). This gene is one of three homologs in a tandem repeat, including Leryth_001243 and Leryth_001244 indicating that tandem gene duplication has expanded this cytochrome P450 subfamily in *L. erythrorhizon* (Fig. S10). Leryth_001243 and Leryth_001244 were not captured in the shikonin subnetwork but their expression is greater in whole root versus above ground tissue (Table S3). Lastly, the phylogeny of sequences homologous to the third cytochrome P450 candidate (Leryth_000257), which encodes a CYP89A2-like enzyme, shows a smaller group of Boraginales sequences without the rounds of expansion present in the other two trees (Fig. S11).

The production of geranylhydroquinone (GHQ) from 3-geranyl-4HBA (Fig. 1) may occur via decarboxylation and subsequent hydroxylation or a single oxidative decarboxylation event^4^. In addition to cytochromes P450, we examined the generated shikonin network for non-cytochrome P450 candidate genes that may function in either hypothesized mechanism. One candidate to consider is a prephenate dehydrogenase-like (PDH-like) gene. PDH catalyzes oxidative decarboxylation of prephenate to 4-hydroxyphenylpyruvate for synthesis of tyrosine^46^. Leryth_001358 encodes a PDH-like protein and is coexpressed with *LePGT1* in the N2 subnetwork (Fig. 5b; Table S7). Similar to other genes in our analysis, the phylogeny of PDHs shows an expansion of this gene family within the Boraginaceae (Fig. S12). Coexpression of the PDH-like gene may simply be related to the connection between shikonin and aromatic amino acid metabolism via phenylpropanoid metabolism, further research is needed to determine if a duplicated PDH could evolve to utilize another 4-hydroxylated substrate.

Given the dozens of shikonin and alkannin derivatives collectively present in the Boraginaceae^47^, we looked for genes in the shikonin subnetwork that may encode tailoring enzymes involved in the synthesis of shikonin derivatives. Recently, two BAHD acyltransferases, shikonin O-acyltransferase (LeSAT1) and alkannin O-acyltransferase (LeAAT1), were discovered to mediate enantiomer-specific acylation in *L. erythrorhizon*^48^. Neither *LeSAT1* nor *LeAAT1* were recovered in the coexpression networks but expression of both is more abundant in at least one shikonin-abundant condition (Table S3). In our N2 shikonin subnetwork (Fig. 5b; Table S7), two additional genes encoding putative transferases (Leryth_012925, Leryth_015823) were recovered. Phylogenetic analysis of the Leryth_015823 transferase and its homologs places Leryth_015823 in a group that contains all Boraginaceae species in our analysis (Fig. S13). In contrast, phylogenetic analysis of Leryth_012925 and its homologs shows Leryth_012925 on a long branch and lacking closely related homologs in other Boraginaceae (Fig. S14), which may make it a potential candidate for a *L. erythrorhizon*-specific shikonin/alkannin tailoring enzyme that is absent in the other shikonin-producing species in our analysis.

### Coexpression network analysis reveals candidates with links to ubiquinone biosynthesis

It has already been demonstrated that *LePGT1* and *LePGT2* evolved via duplication of a primary metabolic prenyltransferase involved in ubiquinone biosynthesis^19^. Given this previously observed connection, the coexpression of a COQ4 ubiquinone biosynthesis-like gene with *LePGT1*, *LeGPPS*, and *LeCYP76B101* in the N2 network appeared remarkable (Fig. 5b; Table S7). Although the precise biochemical function of COQ4 is unknown it is thought to function as a scaffold protein binding proteins and lipids required for efficient ubiquinone biosynthesis^49^. The phylogenetic tree of the COQ4 gene family (Fig. S15) is strikingly similar to that of the ubiquinone prenyltransferase gene family^19^. Both phylogenies contain two subfamilies of Boraginales sequences. One subfamily has shorter branch lengths and contains a single sequence per species, suggesting that this subfamily has retained the ancestral COQ4 ubiquinone biosynthesis activity (Fig. S15). The second Boraginales subfamily has longer branches and shows a radiation of COQ4 paralogs and includes the candidate gene (Leryth_002195; Fig. S15), which was overexpressed in all shikonin-abundant conditions (Table S3). Given the similarities in the precursors and biosynthetic steps in the ubiquinone and shikonin pathways^19^, this COQ4 paralog (Leryth_002195) could fulfill an analogus function and participate in assembling a shikonin biosynthesis metabolon.

In addition to the COQ4-like genes, we also identified two COQ3-like O-methyltransferase genes in the N2 shikonin subnetwork (Leryth_019821 and Leryth_021171). Leryth_019821 was coexpressed with *LeMDR* and Leryth_021171 was coexpressed with *LeGPPS* and *LeCYP76B101* (Fig. 5b). A phylogenetic tree of the COQ3 gene family suggests that the two copies in *L. erythrorhizon* diverged in an ancestor of the Boraginaceae; the Leryth_021171 subfamily contained a sequence from *M. paniculata*, whereas the Leryth_019821 subfamily appears to be unique to shikonin producing species (Fig. S16). The metabolic significance of this network connection remains enigmatic, though it is possible that these enzyme could function in formation of shikonin derivatives.

A final connection to ubiquinone metabolism uncovered in the coexpression analysis was the recovery of a quinoprotein dehydrogenase gene (Leryth_020454) in the largest N4 subnetwork that coexpressed with *LeGPPS*, *LePGT1*, and *LeCYP76B101* (Fig. S4d; Table S9). Leryth_020454 was also significantly overexpressed in all shikonin-abundant conditions (Table S3). Quinoprotein dehydrogenases catalyze the oxidation of glucose to gluconate with concomitant reduction of ubiquinone to ubiquinol^50^. It is conceivable that such an enzyme could function to maintain shikonins and/or pathway intermediates in reduced states to protect the cell. Alternatively, it could function to ensure a pathway intermediate(s) remains in its reduced form. A similar chemical prerequisite is necessary for transmethylation of the 1,4-naphthoquinone ring of demethylphylloquinone in the vitamin K1 pathway^51^. The phylogeny of quinoprotein dehydrogenases shows two copies of this gene in the Boraginaceae (Fig. S17). The clade that contains Leryth_020454 appears unique to shikonin producers and is absent in *M. panticulata* (Fig. S17). Collectively, the analyses provided here suggest there are multiple genes in the shikonin coexpression network that originated from duplication of ubiquinone pathway genes.

## Discussion

In this study, we downregulated expression of *LeGPPS* to explore the connections linking the shikonin pathway with the pathways supplying its metabolic precursors. In doing so, we showed that the recently discovered LeGPPS, an FPPS with evolved GPPS activity^18^, is required for shikonin production (Fig. 2) and that LeGPPS supplies GPP precursor to the shikonin pathway using MVA-pathway derived IPP/DMAPP (Fig. 3). We also performed a series of computational analyses to investigate the evolutionary history of metabolic innovation in the shikonin pathway. Synteny analysis of the *L. erythrorhizon* genome revealed one syntenic block in contigs containing *LeFPPS1* and *LeFPPS3* (Fig. S5) suggesting that WGD in the Boraginaceae was responsible for a duplication giving rise to these canonical FPPS paralogs (Fig. 6). However, the absence of shared synteny between *LeGPPS* and any other *FPPS* genes, suggests that *LeGPPS* did not arise via WGD. There is also no clear evidence of tandem duplication, and the presence of introns likely rules out retro duplication similar to what occurred with *PGT* evolution^19^. Instead, conservation of intron positions between *LeGPPS* and other *FPPS* genes (Fig. S6) is consistent with a segmental or DNA transposition event.

Wisecaver *et al.*^25^ previously showed that network analysis based on abundant coexpression data (*i.e.* hundreds of RNA-seq and/or microarray samples) is a powerful strategy for high-throughput discovery of genes involved in specialized metabolic pathways in plants. We utilized a similar computational approach here with a limited but strategically selected set of transcriptome samples (N=14) to construct a shikonin metabolic network model. We chose to focus our analysis on *LeGPPS*^18^ (Fig. 2), *LePGT1*^19^, *LeCYP76B101*^33, 34^, and *LeMDR*^35^ given their demonstrated roles in shikonin metabolism. Using conventional differential gene expression analysis to refine the gene coexpression matrix, we uncovered a *L. erythrorhizon* shikonin gene network model that predicts strong associations between MVA pathway genes and known shikonin biosynthesis genes, as well as links between shikonin genes and several uncharacterized enzyme-coding genes (Fig. 5) that present new candidates for missing shikonin biosynthesis steps (Fig. 1).

Similar to *LeGPPS* (Fig. 6) and the *PGTs*^19^, Boraginales-specific gene family expansions were observed in the phylogenies (Figs. S8,S9,S11-S17) of the gene candidates identified by coexpression network modeling (Table 1). Therefore, gene duplication appears to be the primary mechanism contributing to metabolic innovation in the Boraginales. Synteny analysis suggests that WGD was unlikely to be responsible for the expansion in the gene families for these candidates (data not shown). Furthermore, examination of the genomic regions surrounding these candidates suggests that tandem duplication did not contribute to their respective gene family expansions either, except for the cytochrome P450 encoded by Leryth_001242 (Fig. S10). Though Leryth_001243 and Leryth_001244 were not candidates identified in the coexpression network, their transcript abundance is higher in roots than in aboveground tissues (Table S3). These cytochromes P450 are predicted to encode CYP76A2-like enzymes. Other CYP76A members have been found to catalyze oxidation cascades involved in formation of terpenoid-derived specialized metabolites^52, 53^, thus making Leryth_001242 and its paralogs intriguing shikonin pathway candidate genes.

The shikonin pathway relies on precursors from both isoprenoid and phenylpropanoid metabolism. Inhibitor experiments with *LeGPPS*-RNAi lines led us to discover an additional layer of regulatory complexity coordinating flux between the phenylpropanoid, MVA, and MEP pathways. Inhibition with the MEP pathway inhibitor fosmidomycin, for example, unexpectedly led to increased shikonin levels in both the *EV*-26 and *LeGPPSi*-45 lines (Fig. 3). This not only provides further evidence that neither IPP/DMAPP derived from the MEP pathway, nor GPP produced from MEP pathway-derived IPP/DMAPP, is exported to the cytoplasm for shikonin biosynthesis but it likely points to an increase in flux through the MVA pathway due to the impairment of the MEP pathway. This therefore implicates the existence of unknown factors modulating distribution of flux from central carbon metabolism between the MVA and MEP pathways, adding another level of control to the complex regulation of the parallel routes in plants^15^.

Comparative RNA-seq analysis of *EV*-26 and *LeGPPSi*-45 hairy root lines revealed that downregulation of *LeGPPS* results in transcriptional changes of genes throughout the terpenoid and phenylpropanoid metabolic networks (Fig. 4). The decreased expression of upstream MVA pathway genes and increased expression of genes encoding cytoplasmic enzymes utilizing IPP/DMAPP (i.e. NUDX1^17^ and FPPS) may indicate that IPP/DMAPP accumulates when the LeGPPS step is limiting. The increased pool of IPP/DMAPP may then be sensed by the cell leading to transcriptional reprogramming of isoprenoid metabolism to redirect the C5 building blocks toward other products. While levels of sterols (cytoplasmic IPP/DMAPP-derived product) and abscisic acid (plastidial IPP/DMAPP-derived product) were not significantly different, the levels of ubiquinones (mitochondrial IPP/DMAPP-derived product) were increased by 36% in *LeGPPSi*-45 lines compared to *EV*-26 lines (Fig. S18). The observed increase in ubiquinone levels is also noteworthy because it further suggests that its precursor pools are shared with the shikonin pathway.

The WRKYs are strong candidates for factors coordinately regulating expression of phenylpropanoid and terpenoid metabolic genes. As one of the largest classes of plant transcription factors, they are involved in regulating processes in response to a number of developmental cues and environmental stimuli. Moreover, they can act as activators or repressors and in doing so they create a regulatory network modulating signaling events from organelles and the cytoplasm to the nucleus^54^. Here, we found that 72 of the 152 genes in the N2 shikonin subnetwork, including *LePGT1, LeGPPS*, *LeMDR*, and *CYP76B101*, contain a canonical W-box element sequence motif (T)TGAC(C/T) (Table S11) recognized by the WRKY family of transcription factors^42^. From our analyses we identified five candidate transcription factors containing WRKY domains in the N2 shikonin subnetwork (Fig. 5b) including two, Leryth_027519 and Leryth_002564, which were both overexpressed in all shikonin-abundant conditions in the analyzed RNA-seq datasets (Table S3).

In addition to sharing 4-HBA and MVA-derived prenyl diphosphate metabolic precursors and having a common origin of their prenyltransferase genes, the shikonin and ubiquinone pathways rely on multiple analogous biochemical ring modifications^19^. This raises the prospect that neofunctionalization of duplicated ubiquinone biosynthesis genes facilitated evolution of the shikonin pathway. Considering this hypothesis, we explored the shikonin coexpression subnetworks for other connections to ubiquinone biosynthesis-like genes. Interestingly, COQ3-like O-methyltransferase and a quinoprotein dehydrogenase genes were found in the coexpression network that are unique to shikonin-producing species (Figs. S16 and S17). Whether these genes function in shikonin metabolism or point to another functional connection between shikonin and ubiquinone remains unclear. Moreover, we identified that in addition to encoding a canonical COQ4, *L. erythrorhizon* has a COQ4-like gene that was coexpressed with *LePGT1*, *LeGPPS*, and *LeCYP76B101* in the N2 network and was overexpressed in shikonin-abundant conditions (Fig. 5b; Tables S3 and S7). COQ4 is a scaffold protein found in plants, fungi, and animals, including humans, that is required for ubiquinone biosynthesis. While its specific function is unknown, it binds proteins and lipids and thus likely assembles a metabolon for efficient ubiquinone biosynthesis^49^. Whether COQ4-like functions similarly in shikonin biosynthesis is an open question that should be explored, especially considering any insight may inform the function of the canonical COQ4 found throughout eukaryotes. Given that shikonin is abundant and non-vital, it may provide a better model for genetically studying the COQ4 gene family.

In summary, our study has i) indicated transcriptional and metabolic connections linking the shikonin pathway with it precursor pathways; ii) established a shikonin coexpression network model that includes genes encoding candidates for missing shikonin pathway steps and regulatory factors; iii) revealed instances of Boraginales-specific gene family expansion facilitated by duplication events for genes in the shikonin metabolic network; and iv) uncovered evolutionary links between shikonin metabolic network genes and ubiquinone pathway genes. The evolution of other plant specialized 1,4-naphthoquinone pathways appears to be linked to primary metabolic quinone pathways^21, 22^. Thus, we expect that the evolutionary mechanistic insights gained here, combined with the demonstration that a robust coexpression network can be built from a small set of RNA-Seq experiments relying on spatial-and condition-specific metabolite correlations, can be used to guide further investigation into the convergent evolution of specialized 1,4-naphthoquinone metabolism in plants.

## Materials and Methods

### Plant materials and hairy root culturing

*Lithospermum erythrorhizon* (accession Siebold & Zucc.) seeds were obtained from the Leibniz Institute of Plant Genetics and Crop Plant Research (IPK) seed bank (Gatersleben, Germany). Propagation of plants to bulk seeds and the generation and maintenance of hairy roots were performed as done previously^19^.

### Generation of *LeGPPSi* and empty-vector control hairy root lines

The *LeGPPS*-RNAi (*LeGPPSi*) construct was created by synthesizing (Genscript, Piscataway, NJ) spliced fragments of the *LeGPPS* coding region corresponding to nucleotides 165-727 and 165-519, the latter in antisense orientation to create a hairpin structure. A 5’-CACC sequence was added for subcloning into pENTR^TM^/D-TOPO (Invitrogen^TM^, Carlsbad, CA) and subsequent transfer into the destination vector, pB2GW7^55^, by recombination using LR Clonase Enzyme Mix^TM^ (Invitrogen). The final construct, pB2GW7*-GPPSi*, was transformed into *Agrobacterium rhizogenes* strain ATCC 15834 competent cells by freeze-thaw transformation^56^ and plated on Nutrient Broth (NB) agar containing 50 µg/mL spectinomycin for selection.

*L. erythrorhizon* hairy root *GPPSi* lines were generated by applying prepared cultures of *A. rhizogenes* containing the pB2GW7*-GPPSi* construct to wounded stems of *L. erythrorhizon* plants in tissue culture as previously described ^19^. Emergent roots from plants 2-4 weeks after infection were excised and transferred to Gamborg B5 media plates containing 3% sucrose and 200 µg/mL cefotaxime to eliminate *A*. *rhizogenes.* After 2 weeks, hairy roots were transferred to Gamborg B5 media containing 3% sucrose and 10 mg/L Basta for selection for 2 weeks. Hairy root lines transformed by *A. rhizogenes* carrying an empty pB2GW7 vector were generated in parallel as controls.

### RNA extraction and qRT-PCR analysis

Total RNA was extracted from ∼100 mg of flash-frozen hairy root tissue and qRT-PCR reactions were performed using a QuantStudio^TM^ 6 (ThermoFisher) as previously described ^19^. Expression of *LeGPPS* and *LeGPPS2* was measured with comparative quantification using the 2^−ΔΔCT^ method ^57^. Primers were designed using Primer-BLAST on NCBI ^58^ (Table S15). Expression was normalized to *L. erythrorhizon* glyceraldehyde 3-phosphate dehydrogenase (*LeGAPDH*) ^59^.

### Metabolite extraction and quantification

Extraction and analysis of ABA by liquid chromatography coupled with tandem mass spectrometry (LC-MS/MS) was performed as previously described^60^. Extraction of total shikonins from growth media of hairy root cultures and quantification on an Agilent 1260 Infinity high performance liquid chromatography with diode array detection (HPLC-DAD) system (Agilent Technologies) was done as previously described ^19^. Sterols were extracted from 100-200 mg of ground flash-frozen hairy root tissue, derivatized with BSTFA, and analyzed on an Agilent 7890B gas chromatograph (GC) coupled with a 5977A mass spectrometer (MS) equipped with a DB-5MS column (30 m × 0.25 mm × 0.25 μm film; Agilent Technologies) and employing Chemstation software as previously described^16^. Ubiquinones were extracted from 100-200 mg of ground flash-frozen fresh tissue in 3 mL of 95% ethanol spiked with 4 nmol ubiquinone-4 internal standard and incubated overnight with shaking at 4°C. The next day, samples were centrifuged at 500 x *g* to pellet debris. Then, 1.5 mL of water was added to supernatant and partitioned twice with 4.5 mL hexane. The hexane layers were combined and concentrated under nitrogen gas at 37°C. Nearly dry samples were resuspended in 1 ml 90:10 methanol:dichloromethane and filtered through 0.2 µm PTFE syringe filters. Care was taken throughout the extraction process to protect samples from light. Samples were analyzed by HPLC-DAD on an Agilent Zorbax SB-C18 column (5 µm, 250 x 4.6 mm) thermostatted at 25°C and eluted in isocratic mode with 30% 60:40 isopropanol:hexanes and 70% 80:20 methanol:hexanes^8^. Ubiquinones were detected spectrophotometrically at 255 nm and had retention times of 4.8 min for ubiquinone-4, 11.4 min for ubiquinone-9, and 14.3 min for ubiquinone-10. Instrument operation and data analysis steps were performed through the Agilent ChemStation software. Quantification of ubiquinones was done by DAD using signals obtained in the linear range of calibration standards (0.0313, 0.0625, 0.125, 0.250 and 0.500 nmol). The data were corrected for recovery according to the ubiquinone-4 internal standard, and final quantifications were made using linear regression. Differences in total shikonin and ubiquinone-9 and ubiquinone-10 content produced by empty-vector control and *LeGPPSi* lines (n = 4 biological replicates) were analyzed using one-way ANOVA and means were compared with Tukey’s HSD post-hoc test at a 95% significance level.

### RNA-sequencing analysis of *LeGPPSi* and empty-vector control lines

For RNA-seq analysis of *L. erythrorhizon EV*-26 and *LeGPPSi*-45, three independent hairy root cultures of each line were started in liquid Gamborg B5 media containing 3% sucrose and grown at 28°C in 100 µE m^−2^s^−1^ light. After two weeks, the hairy roots were transferred to M9 media containing 3% sucrose and darkness for six days. The hairy roots were then frozen in liquid nitrogen, ground by mortar and pestle, and RNA was extracted from ∼100 mg of tissue as described above. Library construction (NEBNext Ultra RNA Library Prep Kit, New England Biolabs Inc.) from 1 µg RNA, Illumina sequencing, and analyses of DEGs were performed by Novogene Corporation Inc. (Sacramento, CA). Paired-end clean reads were mapped to the *L. erythrorhizon* reference genome^19^ using HISAT2 software^61^ For each sequenced library, read counts were adjusted by TMM^62^ and DEG analysis was performed using DESeq2^63^ with *p*-value adjusted using an FDR calculated with Benjamini–Hochberg (BH) methods^64^. Genes were considered significantly differentially expressed if they had a BH-adjusted *p*-value of 0.005 and a log2 fold change of 1. Kyoto Encyclopedia of Genes and Genomes (KEGG) pathway enrichment analyses of DEGs was implemented by the clusterProfiler R package^65^ and KEGG pathways with BH-adjusted *p*-value < 0.05 were considered significantly enriched. The raw data were submitted to the Sequence Read Archive (http://www.ncbi.nlm.nih.gov/sra/) and are available at the NCBI Sequence Read Archive (PRJNAXXXXXX).

### Analysis of transcriptomes used to build shikonin gene coexpression networks

Illumina RNA-seq reads of *L. erythrorhizon* root periderm, root vascular, and hairy root cultures were generated as described^19^ and are available at the NCBI Sequence Read Archive (PRJNA596998). Additional Illumina RNA-seq reads of *L. erythrorhizon* whole roots and above ground tissue (pooled leaves and stems) were downloaded from the NCBI SRA database, experiments SRR3957230 and SRR3957231 respectively. *L. erythrorhizon* gene functional annotations were downloaded from^19^.

*L. erythrorhizon* RNA-seq raw reads were error corrected using the Tadpole (default parameters; software last modified June 27, 2017) program from the BBMap software package (https://sourceforge.net/projects/bbmap/). Gene expression was quantified with Kallisto^66^ by aligning the error corrected reads to a collection of the longest transcript per gene of the *L. erythrorhizon* genome. The *LePS-2* gene was previously implicated in shikonin biosynthesis ^27^ but was not present in v1.0 of the *L. erythrorhizon* gene set^19^. Therefore, we identified a putative coding sequence for *LePS-2* in the *L. erythrorhizon* genome assembly manually and added its sequence to the total gene set prior to gene expression quantification.

Analyses of differential gene expression was performed using the edgeR package^67^. Gene expression counts were normalized using the TMM (trimmed mean of M values) method^62^. Exact tests were conducted using a trended dispersion value and a double tail reject region. FDRs were calculated using the BH procedure^64^. Genes that did not have a significant differential expression status in at least one comparison (FDR < 0.1) were excluded from downstream coexpression analyses.

### Coexpression network analysis

Raw gene expression counts were normalized using the transcripts per million method and transformed using the variance-stabilizing transformation method in DESeq2^63^, and global gene coexpression networks were constructed as previously described^25^. Briefly, a Pearson’s correlation coefficient (PCC) was calculated between gene pairs and converted into a mutual rank (MR) using scripts available for download on GitHub (https://github.rcac.purdue.edu/jwisecav/coexp-pipe). MR scores were transformed to network edge weights using the exponential decay function 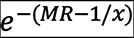; four different networks were constructed with *x* set to 5, 10, 25, and 50, respectively. Edges with a weight < 0.01 were trimmed from the global network. Modules of coexpressed genes were detected using ClusterOne v1.0 using default parameters^31^. Overlapping modules within each coexpression network were combined by collapsing all modules containing the known Shikonin pathway genes *LeGPPS*^18^, *LePGT1*^13^, *LeCYP76B101*^34^ and *LeMDR*^68^ into a subnetwork. Modules were visualized in Cytoscape using the spring embedded layout. Tests for functional enrichment of Gene Ontology (GO) terms in the different shikonin subnetworks (Table S10) were performed using hypergeometric tests using the SciPy library hypergeom, and p-values were adjusted for multiple comparisons using the StatsModels library multitest using the BH procedure^64^. GO terms and other gene functional annotations were taken from^19^.

### Promoter analysis

Nucleic acid sequence motifs enriched in promoter regions of genes in the N2 shikonin subnetwork (N=152) were identified with Motif Indexer^41^ using a 1000 base pair window upstream of all transcriptional start sites using the same upstream region of all *L. erythrorhizon* genes as background. Identified motifs were consolidated and ranked using the KeyMotifs.pl perl script provided by Motif Indexer. To calculate a false discovery rate, 1000 random sets of 152 genes were run through Motif Indexer determine a *p*-value threshold. No motif identified from a random gene set had a *p*-value less than 1×10^-9^.

### Phylogenetic analysis

To construct gene phylogenies, the gene family containing the best *Arabidopsis thaliana* BLAST hit to the query gene was downloaded from the PLAZA 4.0 Dicots comparative genomics database^43^. Homology between the predicted proteomes of *L. erythrorhizon* and 18 additional Borginales was determined with OrthoFinder v2.1.2 using the following parameters: -S diamond -M msa -T fasttree^69^. OrthoFinder orthogroups containing the query gene were combined with the Plaza 4.0 gene family to obtain the final sequence sets. Sequences were aligned with MAFFT^70^ using the E-INS-I strategy and following parameters: --maxiterate 1000 --bl 45 --op 1.0 --retree 3. The maximum likelihood phylogeny was constructed using IQ-TREE^71^ using the built in ModelFinder to determine the best-fit substitution model^72^ and performing SH-aLRT and the ultrafast bootstrapping analyses with 1000 replicates each. For the cytochrome P450 and acetyltransferase gene candidates, because the PLAZA 4.0 gene families were so large, a quick guide tree of the entire gene family was built using FastTree^73^. Regions of the guide tree that contained candidate genes of interest were identified; sequences within these regions were realigned using MAFFT, and phylogenies were built using IQ-TREE as described above.

### Synteny analysis

Regions of shared synteny within the genome of *L. erythrorhizon* were detected using SynMap2 on the online Comparative Genomics Platform (CoGe) using default settings with the exception that the merge syntenic blocks algorithm was set to Quota Align Merge, syntenic depth algorithm was set to Quota Align, and the CodeML option was activated to calculate substitution rates between syntenic CDS pairs. For syntenic blocks containing genes of interest and their homologs, the encompassing contigs were aligned using promer of the MUMmer4 alignment system^74^.

## Data availability

Raw sequencing reads have been deposited in the Sequence Read Archive database under access number PRJNAXXXXXX.

## Acknowledgements

We dedicate this manuscript in memory of our late colleague Dr. Chunhua Zhang whose passion and curiosity for science served as an inspiration to us all. This work was supported by start-up funds from Purdue University to S.A.M.M., J.H.W. and J.R.W.; the NSF Dimensions of Biodiversity Program under Grant No. DEB-1831493 to J.H.W.; and by a Fellowship from the Anandamahidol Foundation (Thailand) to T.S. This work was supported by the USDA National Institute of Food and Agriculture Hatch Project numbers 177845 to J.R.W. and 1016057 to J.H.W. We also want to acknowledge use of the Metabolite Profiling Facility at the Bindley Bioscience Center, a core facility of the NIH-funded Indiana Clinical and Translational Sciences Institute.

## Supporting information figure legends

**Fig. S1**

**Effect of *LeGPPS* RNAi downregulation on expression of shikonin pathway genes.** The average log_2_fold-change in expression for each gene in *LeGPPSi*-45 lines compared to *EV-26* lines. The CYP76B101 gene (*) was not included in this DE analysis as the non stranded RNAseq library was unable to distinguish between CYP76B101 (Leryth_016594) and Leryth_016593, which occupies the same genomic location but is encoded on the opposite strand. See Fig. 1 legend for abbreviations and Table S4 for gene descriptions.

**Fig. S2**

**Kyoto Encyclopedia of Genes and Genomes (KEGG) term enrichment analysis of genes downregulated in *LeGPPSi*-45 compared to *EV*-26 hairy root lines.** KEGG pathways with corrected *p*-value < 0.05 were considered significantly enriched by differential expressed genes. *EV*-26, empty-vector control line 26; *LeGPPSi*-45, *LeGPPS*-RNAi line 45.

**Fig. S3**

**Exogenous application of 4-hydroxybenzoate (4-HBA) does not restore shikonin production in *LeGPPSi*-45 lines.** All data are means ± SEM (n = 4 biological replicates). Different letters indicate significant differences via analysis of variance (ANOVA) followed by post-hoc Tukey test (α= 0.05).

**Fig. S4a**

**Shikonin subnetwork N1.** Network map of genes coexpressed with *LePGT*, *LeGPPS*, *LeCYP76B101*, and *LeMDR* using the N1 global coexpression network. Nodes are colored according to the gene’s coexpression status with known shikonin genes. Network maps were drawn using a Fruchterman-Reingold force-directed layout using the edge-weighted spring embedded layout in cytoscape (https://cytoscape.org).

**Fig. S4b Shikonin subnetwork N2.** Network map of genes coexpressed with *LePGT*, *LeGPPS*, *LeCYP76B101*, and *LeMDR* using the N2 global coexpression network.

Network maps are drawn as described in Supplemental Figure S1a.

**Fig. S4c Shikonin subnetwork N3.** Network map of genes coexpressed with *LePGT*, *LeGPPS*, *LeCYP76B101*, and *LeMDR* using the N3 global coexpression network.

Network maps are drawn as described in Supplemental Figure S1a.

**Fig. S4d Shikonin subnetwork N4.** Network map of genes coexpressed with *LePGT*, *LeGPPS*, *LeCYP76B101*, and *LeMDR* using the N4 global coexpression network.

Network maps are drawn as described in Supplemental Figure S1a.

**Fig. S5**

**Shared synteny between FPPS homologs.** Mummerplot of syntenic block containing farnesyl pyrophosphate synthase homologs Leryth_005102 (*LeFPPS1*) and Leryth_010152 *(LeFPPS3*). The locations of the homologs are indicated by the red arrows.

**Fig. S6**

**Conservation of intron locations in the *FPPS* gene family.** Multiple sequence alignment showing conservation of intron locations (indicated by xxxxxxxxxx) between *LeGPPS* (Leryth_023382), *LeFPPS1* (Leryth_005102), *LeFPPS2* (Leryth_007856), and *LeFPPS3* (Leryth_010152).

**Fig. S7**

**Maximum likelihood phylogeny of FPPS gene family in green plants.** Nodes with IQ-TREE support values > 95 are indicated by numbers on the preceding branch. The branches and outer color bar are color-coded to match the taxonomic classification of each sequence. The tree is rooted on Chlorophytes. Non-Boraginales sequences containing a Histidine residue adjacent to the conserved Asp-rich motif are indicated by an asterisk (Fv = F. vesca, Pb = P. bretschneideri, St = S. tuberosum).

**Fig. S8**

**Maximum likelihood phylogeny of cytochrome CYP76B genes and homologs in green plants.** Nodes with IQ-TREE support values > 95 are indicated by grey circles on the preceding branch. The branches and outer color bar are color-coded to match the taxonomic classification of each sequence. The tree is rooted based on rough guide tree of entire P450 gene family (Plaza Dicots 4.0 HOM04D000003).

**Fig. S9**

**Maximum likelihood phylogeny of cytochrome P450 candidate Leryth_001242 and homologs in green plants.** Nodes with IQ-TREE support values > 95 are indicated by numbers on the preceding branch. The branches and outer color bar are color-coded to match the taxonomic classification of each sequence. The tree is rooted based on rough guide tree of entire cytochrome P450 gene family (Plaza Dicots 4.0 HOM04D000003).

**Fig. S10**

**UCSC Genome Browser region for P450 candidate Leryth_001242.**

**Fig. S11**

**Maximum likelihood phylogeny of cytochrome P450 candidate Leryth_000257 and homologs in green plants.** Nodes with IQ-TREE support values > 95 are indicated by grey circles on the preceding branch. The branches and outer color bar are color-coded to match the taxonomic classification of each sequence. Tree is midpoint rooted. The tree is rooted based on rough guide tree of entire cytochrome P450 gene family (Plaza Dicots 4.0 HOM04D000435).

**Fig. S12**

**Maximum likelihood phylogeny of the prephenate dehydrogenase gene family in green plants.** Nodes with IQ-TREE support values > 95 are indicated by numbers on the preceding branch. The branches and outer color bar are color-coded to match the taxonomic classification of each sequence. The tree is rooted on Chlorophytes.

**Fig. S13**

**Maximum likelihood phylogeny of AT candidate Leryth_015823 and homologs in green plants.** Nodes with IQ-TREE support values > 95 are indicated by numbers on the preceding branch. The branches and outer color bar are color-coded to match the taxonomic classification of each sequence. The tree is rooted based on rough guide tree of entire AT gene family (Plaza Dicots 4.0 HOM04D000075).

**Fig. S14**

**Maximum likelihood phylogeny of AT Candidate Leryth_012925 and homologs in green plants.** Nodes with IQ-TREE support values > 95 are indicated by numbers on the preceding branch. The branches and outer color bar are color-coded to match the taxonomic classification of each sequence. The tree is rooted based on rough guide tree of entire AT gene family (Plaza Dicots 4.0 HOM04D000339).

**Fig. S15**

**Maximum likelihood phylogeny of COQ4 gene family in green plants.** Nodes with IQ-TREE support values > 95 are indicated by numbers on the preceding branch. The branches and outer color bar are color-coded to match the taxonomic classification of each sequence. The tree is rooted on Chlorophytes.

**Fig. S16**

**Maximum likelihood phylogeny of the COQ3 gene family in green plants.** Nodes with IQ-TREE support values > 95 are indicated by numbers on the preceding branch. The branches and outer color bar are color-coded to match the taxonomic classification of each sequence. The tree is rooted on Chlorophytes.

**Fig. S17**

**Maximum likelihood phylogeny of quinoprotein dehydrogenase gene family in green plants.** Nodes with IQ-TREE support values > 95 are indicated by numbers on the preceding branch. The branches and outer color bar are color-coded to match the taxonomic classification of each sequence. The tree is rooted on mosses.

**Fig. S18**

**Pool sizes of sterols (a), ubiquinones (b), and abscisic acid (ABA) (c) measured in empty-vector control line 26 (EV-26) and LeGPPS RNAi line 45 (LeGPPSi-45).**

Metabolites were measure at 6 d after transfer of 14-d-old hairy roots to M9 and darkness. All data are means ± SEM (n = 3–4 biological replicates). Statistically significant differences are indicated (*P < 0.05, Student’s t test).

## Supporting information table legends

**Table S1**

Differentially expressed genes in LeGPPSi-45 compared to EV-26 Lithospermum erythrorhizon hairy roots

**Table S2**

Summary of differentially expressed genes in publicly available comparative RNA-seq experiments

**Table S3**

Differential expression analyses

**Table S4**

List of known and candidate shikonin proteins

**Table S5**

Summary of network module identification using ClusterOne

**Table S6**

N1 shikonin subnetwork genes

**Table S7**

N2 shikonin subnetwork genes

**Table S8**

N3 shikonin subnetwork genes

**Table S9**

N4 shikonin subnetwork genes

**Table S10**

Gene Ontology functional enrichment analysis

**Table S11**

Motif enrichment results

**Table S12**

Gene families used to construct phylogenies

**Table S13**

Boraginales species used in OrthoFinder analysis

**Table S14**

CodeML substitution rates between syntenic CDS pairs.

**Table S15**

Primers used in this study

## Supporting information dataset legends

**Dataset S1**

**Network modules.** Each line corresponds to a module and contain the network (N1-N4), module number, size, density, total weight of edges internal to and exiting a module, the value of the quality function, a P-value and the list of genes found within each module. Columns are separated by commas.

## REFERENCES

1 Papageorgiou VP, Assimopoulou AN, Couladouros EA, Hepworth D, Nicolaou KC. The chemistry and biology of alkannin, shikonin, and related naphthazarin natural products. Angew Chemie-Int Ed 1999; 38: 270–300.

2 Skoneczny D, Weston P, Zhu X, Gurr G, Callaway R, Barrow R et al. Metabolic Profiling and Identification of Shikonins in Root Periderm of Two Invasive Echium spp. Weeds in Australia. Molecules 2017; 22: 330.

3 Zhu X, Skoneczny D, Weidenhamer JD, Mwendwa JM, Weston PA, Gurr GM et al. Identification and localization of bioactive naphthoquinones in the roots and rhizosphere of Paterson’s curse (Echium plantagineum), a noxious invader. J Exp Bot 2016; 67: 3777–3788.

4 Widhalm JR, Rhodes D. Biosynthesis and molecular actions of specialized 1,4-naphthoquinone natural products produced by horticultural plants. Hortic Res 2016; 3: 16046.

5 Wang F, Yao X, Zhang Y, Tang J. Synthesis, biological function and evaluation of Shikonin in cancer therapy. Fitoterapia 2019; 134: 329–339.

6 Schmid HV, Zenk MH. p-hydroxybenzoic acid and mevalonic acid as precursors of the plant naphthoquinone alkannin. Tetrahedron Lett 1971; 12: 4151–4155.

7 Loscher R, Heide L. Biosynthesis of p-Hydroxybenzoate from p-Coumarate and p-Coumaroyl-Coenzyme A in Cell-Free Extracts of Lithospermum erythrorhizon Cell Cultures. Plant Physiol 1994; 106: 271–279.

8 Block A, Widhalm JR, Fatihi A, Cahoon RE, Wamboldt Y, Elowsky C, et al. The Origin and Biosynthesis of the Benzenoid Moiety of Ubiquinone (Coenzyme Q) in Arabidopsis. Plant Cell 2014; 26: 1938–1948.

9 Soubeyrand E, Johnson TS, Latimer S, Block A, Kim J, Colquhoun TA et al. The Peroxidative Cleavage of Kaempferol Contributes to the Biosynthesis of the Benzenoid Moiety of Ubiquinone in Plants. Plant Cell 2018; 30: 2910–2921.

10 Yazaki K, Kataoka M, Honda G, Severin K, Heide L. cDNA Cloning and Gene Expression of Phenylalanine Ammonia-Lyase in Lithospermum erythrorhizon. Biosci Biotechnol Biochem 1997; 61: 1995–2003.

11 Yamamura Y, Ogihara Y, Mizukami H. Cinnamic acid 4-hydroxylase from Lithospermum erythrorhizon: cDNA cloning and gene expression. Plant Cell Rep 2001; 20: 655–662.

12 Singh RS, Gara RK, Bhardwaj PK, Kaachra A, Malik S, Kumar R et al. Expression of 3-hydroxy-3-methylglutaryl-CoA reductase, p-hydroxybenzoate-m-geranyltransferase and genes of phenylpropanoid pathway exhibits positive correlation with shikonins content in arnebia [Arnebia euchroma (Royle) Johnston]. BMC Mol Biol 2010; 11: 1471–2199.

13 Yazaki K, Kunihisa M, Fujisaki T, Sato F. Geranyl Diphosphate:4-Hydroxybenzoate Geranyltransferase fromLithospermum erythrorhizon: Cloning and characterization of a key enzyme in shikonin biosynthesis. J Biol Chem 2002; 277: 6240–6246.

14 Yazaki K, Fukui H, Tabata M. Isolation of the intermediates and related metabolites of shikonin biosynthesis from Lithospermum erythrorhizon cell cultures. Chem Pharm Bull (Tokyo) 1986; 34: 2290–2293.

15 Vranová E, Coman D, Gruissem W. Network analysis of the MVA and MEP pathways for isoprenoid synthesis. Annu Rev Plant Biol 2013; 64: 665–700.

16 Henry LK, Gutensohn M, Thomas ST, Noel JP, Dudareva N. Orthologs of the archaeal isopentenyl phosphate kinase regulate terpenoid production in plants. Proc Natl Acad Sci 2015; 112: 10050–10055.

17 Henry LK, Thomas ST, Widhalm JR, Lynch JH, Davis TC, Kessler SA et al. Contribution of isopentenyl phosphate to plant terpenoid metabolism. Nat Plants 2018; 4: 721–729.

18 Ueoka H, Sasaki K, Miyawaki T, Ichino T, Tatsumi K, Suzuki S et al. A Cytosol-Localized Geranyl Diphosphate Synthase from Lithospermum erythrorhizon and Its Molecular Evolution. Plant Physiol 2020; 182: 1933–1945.

19 Auber RP, Suttiyut T, McCoy RM, Ghaste M, Crook JW, Pendleton AL et al. Hybrid de novo genome assembly of red gromwell (Lithospermum erythrorhizon) reveals evolutionary insight into shikonin biosynthesis. Hortic Res 2020; 7: 82.

20 Tang CY, Li S, Wang YT, Wang X. Comparative genome/transcriptome analysis probes Boraginales’ phylogenetic position, WGDs in Boraginales, and key enzyme genes in the alkannin/shikonin core pathway. Mol Ecol Resour 2020; 20: 228–241.

21 Meyer GW, Bahamon Naranjo MA, Widhalm JR. Convergent evolution of plant specialized 1,4-naphthoquinones: metabolism, trafficking, and resistance to their allelopathic effects. J Exp Bot 2021; 72: 167–176.

22 McCoy RM, Utturkar SM, Crook JW, Thimmapuram J, Widhalm JR. The origin and biosynthesis of the naphthalenoid moiety of juglone in black walnut. Hortic Res 2018; 5: 67.

23 Tholl D. Biosynthesis and Biological Functions of Terpenoids in Plants. In: Advances in Biochemical Engineering/Biotechnology. 2015, pp 63–106.

24 Usadel B, Obayashi T, Mutwil M, Giorgi FM, Bassel GW, Tanimoto M et al. Co-expression tools for plant biology: opportunities for hypothesis generation and caveats. Plant Cell Environ 2009; 32: 1633–1651.

25 Wisecaver JH, Borowsky AT, Tzin V, Jander G, Kliebenstein DJ, Rokas A. A Global Coexpression Network Approach for Connecting Genes to Specialized Metabolic Pathways in Plants. Plant Cell 2017; 29: 944–959.

26 Song W, Zhuang Y, Liu T. CYP82AR Subfamily Proteins Catalyze C-1′ Hydroxylations of Deoxyshikonin in the Biosynthesis of Shikonin and Alkannin. Org Lett 2021; 23: 2455–2459.

27 Yamamura Y, Sahin FP, Nagatsu A, Mizukami H. Molecular cloning and characterization of a cDNA encoding a novel apoplastic protein preferentially expressed in a shikonin-producing callus strain of Lithospermum erythrorhizon. Plant Cell Physiol 2003; 44: 437–46.

28 Zhao H, Baloch SK, Kong LR, Zhang WJ, Zou AL, Wang XM et al. Molecular cloning, characterization, and expression analysis of LeMYB1 from Lithospermum erythrorhizon. Biol Plant 2014; 58: 436–444.

29 Obayashi T, Kinoshita K. Rank of correlation coefficient as a comparable measure for biological significance of gene coexpression. DNA Res 2009; 16: 249–260.

30 Liesecke F, Daudu D, De Bernonville RD, Besseau S, Clastre M, Courdavault V et al. Ranking genome-wide correlation measurements improves microarray and RNA-seq based global and targeted co-expression networks. Sci Rep 2018. doi:10.1038/s41598-018-29077-3.

31 Wu H, Gao L, Dong J, Yang X. Detecting overlapping protein complexes by rough-fuzzy clustering in protein-protein interaction networks. PLoS One 2014; 9: 471–472.

32 van Dongen S, Abreu-Goodger C. Using MCL to Extract Clusters from Networks. In: *Bacterial Molecular Networks*. Springer, New York, NY, 2012, pp 281–295.

33 Song W, Zhuang Y, Liu T. Potential role of two cytochrome P450s obtained from Lithospermum erythrorhizon in catalyzing the oxidation of geranylhydroquinone during Shikonin biosynthesis. Phytochemistry 2020; 175: 112375.

34 Wang S, Wang R, Liu T, Lv C, Liang J, Kang C et al. CYP76B74 catalyzes the 3’’-hydroxylation of geranylhydroquinone in shikonin biosynthesis. Plant Physiol 2019; 179: 402–414.

35 Zhu Y, Chu S-J, Luo Y-L, Fu J-Y, Tang C-Y, Lu G-H, et al. Involvement of LeMRP, an ATP-binding cassette transporter, in shikonin transport and biosynthesis in Lithospermum erythrorhizon. Plant Biol 2017; 17: 1–9.

36 Rai A, Nakaya T, Shimizu Y, Rai M, Nakamura M, Suzuki H, et al. De Novo Transcriptome Assembly and Characterization of Lithospermum officinale to Discover Putative Genes Involved in Specialized Metabolites Biosynthesis *. 2018.

37 Wu FY, Tang CY, Guo YM, Bian ZW, Fu JY, Lu GH et al. Transcriptome analysis explores genes related to shikonin biosynthesis in Lithospermeae plants and provides insights into Boraginales’ evolutionary history. Sci Rep 2017. doi:10.1038/s41598-017-04750-1.

38 Fang R, Wu F, Zou A, Zhu Y, Zhao H, Zhao H et al. Transgenic analysis reveals LeACS-1 as a positive regulator of ethylene-induced shikonin biosynthesis in Lithospermum erythrorhizon hairy roots. Plant Mol Biol 2016. doi:10.1007/s11103-015-0421-z.

39 Zhang WJ, Su J, Tan MY, Liu GL, Pang YJ, Shen HG et al. Expression analysis of shikonin-biosynthetic genes in response to M9 medium and light in Lithospermum erythrorhizon cell cultures. Plant Cell Tissue Organ Cult 2010; 101: 135–142.

40 Tatsumi K, Yano M, Kaminade K, Sugiyama A, Sato M, Toyooka K et al. Characterization of Shikonin Derivative Secretion in Lithospermum erythrorhizon Hairy Roots as a Model of Lipid-Soluble Metabolite Secretion from Plants. Front Plant Sci 2016; 7: 1–11.

41 Ma S, Bachan S, Porto M, Bohnert HJ, Snyder M, Dinesh-Kumar SP. Discovery of stress responsive DNA regulatory motifs in arabidopsis. PLoS One 2012; 7. doi:10.1371/journal.pone.0043198.

42 Rushton PJ, Torres JT, Parniske M, Wernert P, Hahlbrock K, Somssich IE. Interaction of elicitor-induced DNA-binding proteins with elicitor response elements in the promoters of parsley PR1 genes. EMBO J 1996; 15: 5690–5700.

43 Van Bel M, Diels T, Vancaester E, Kreft L, Botzki A, Van De Peer Y et al. PLAZA 4.0: An integrative resource for functional, evolutionary and comparative plant genomics. Nucleic Acids Res 2018; 46: D1190–D1196.

44 Leebens-Mack JH, Barker MS, Carpenter EJ, Deyholos MK, Gitzendanner MA, Graham SW et al. One thousand plant transcriptomes and the phylogenomics of green plants. Nature 2019; 574: 679–685.

45 Moore BM, Wang P, Fan P, Leong B, Schenck CA, Lloyd JP et al. Robust predictions of specialized metabolism genes through machine learning. Proc Natl Acad Sci U S A 2019; 116: 2344–2353.

46 Maeda H, Dudareva N. The shikimate pathway and aromatic amino Acid biosynthesis in plants. Annu Rev Plant Biol 2012; 63: 73–105.

47 Papageorgiou V, Assimopoulou A, Samanidou V, Papadoyannis I. Recent Advances in Chemistry, Biology and Biotechnology of Alkannins and Shikonins. Curr Org Chem 2006; 10: 2123–2142.

48 Oshikiri H, Watanabe B, Yamamoto H, Yazaki K, Takanashi K. Two BAHD acyltransferases catalyze the last step in the shikonin/alkannin biosynthetic pathway. Plant Physiol 2020; 184: pp.00207.2020.

49 Stefely J. A., D. J. P. Biochemistry of mitochondrial coenzyme Q biosynthesis. Trends Biochem Sci 2017; 42: 824–843.

50 Oubrie A, Rozeboom HJ, Kalk KH, Olsthoorn AJJ, Duine JA, Dijkstra BW. Structure and mechanism of soluble quinoprotein glucose dehydrogenase. EMBO J 1999; 18: 5187–5194.

51 Fatihi A, Latimer S, Schmollinger S, Block A, Dussault PH, Vermaas WFJ et al. A Dedicated Type II NADPH Dehydrogenase Performs the Penultimate Step in the Biosynthesis of Vitamin K1 in Synechocystis and Arabidopsis. Plant Cell 2015; 27: 1730–1741.

52 Guo J, Zhou YJ, Hillwig ML, Shen Y, Yang L, Wang Y et al. CYP76AH1 catalyzes turnover of miltiradiene in tanshinones biosynthesis and enables heterologous production of ferruginol in yeasts. Proc Natl Acad Sci U S A 2013; 110: 12108–12113.

53 Miettinen K, Dong L, Navrot N, Schneider T, Burlat V, Pollier J et al. The seco-iridoid pathway from Catharanthus roseus. Nat Commun 2014; 5. doi:10.1038/ncomms4606.

54 Bakshi M, Oelmüller R. Wrky transcription factors jack of many trades in plants. Plant Signal Behav 2014; 9: 1–18.

55 Karimi M, Inzé D, Depicker A. GATEWAY^TM^ vectors for Agrobacterium-mediated plant transformation. Trends Plant Sci 2002; 7: 193–195.

56 Cui W, Liu W, Wu G. A simple method for the transformation of Agrobacterium tumefaciens by foreign DNA. Chin J Biotechnol 1995; 11: 267–274.

57 Livak KJ, Schmittgen TD. Analysis of relative gene expression data using real-time quantitative PCR and the 2-ΔΔCT method. Methods 2001; 25: 402–408.

58 Ye J, Coulouris G, Zaretskaya I, Cutcutache I, Rozen S, Madden TL. Primer-BLAST: a tool to design target-specific primers for polymerase chain reaction. BMC Bioinformatics 2012; 13: 134.

59 Zhao H, Chang QS, Zhang DX, Fang RJ, Zhao H, Wu FY et al. Overexpression of LeMYB1 enhances shikonin formation by up-regulating key shikonin biosynthesis-related genes in Lithospermum erythrorhizon. Biol Plant 2015; 59: 429–435.

60 McAdam SAM, Brodribb TJ. Mesophyll cells are the main site of abscisic acid biosynthesis in water-stressed leaves. Plant Physiol 2018; 177: 911–917.

61 Kim D, Paggi JM, Park C, Bennett C, Salzberg SL. Graph-based genome alignment and genotyping with HISAT2 and HISAT-genotype. Nat Biotechnol 2019; 37: 907–915.

62 Robinson MD, Oshlack A. A scaling normalization method for differential expression analysis of RNA-seq data. Genome Biol 2010; 11. doi:10.1186/gb-2010-11-3-r25.

63 Love MI, Huber W, Anders S. Moderated estimation of fold change and dispersion for RNA-seq data with DESeq2. 2014; : 1–21.

64 Benjamini Y, Hochberg Y. Controlling the False Discovery Rate: A Practical and Powerful Approach to Multiple Testing. J R Stat Soc Ser B 1995; 57: 289–300.

65 Yu G, Wang L-G, Han Y, He Q-Y. clusterProfiler: an R Package for Comparing Biological Themes Among Gene Clusters. Omi A J Integr Biol 2012; 16: 284–287.

66 Bray NL, Pimentel H, Melsted P, Pachter L. Near-optimal probabilistic RNA-seq quantification. Nat Biotechnol 2016; 34: 525–527.

67 Robinson MD, McCarthy DJ, Smyth GK. edgeR: A Bioconductor package for differential expression analysis of digital gene expression data. Bioinformatics 2009; 26: 139–140.

68 Zhu Y, Lu GH, Bian ZW, Wu FY, Pang YJ, Wang XM et al. Involvement of LeMDR, an ATP-binding cassette protein gene, in shikonin transport and biosynthesis in Lithospermum erythrorhizon. BMC Plant Biol 2017; 17: 1–10.

69 Emms DM, Kelly S. OrthoFinder: solving fundamental biases in whole genome comparisons dramatically improves orthogroup inference accuracy. Genome Biol 2015; 16: 1–14.

70 Katoh K, Standley DM. MAFFT multiple sequence alignment software version 7: Improvements in performance and usability. Mol Biol Evol 2013; 30: 772–780.

71 Nguyen LT, Schmidt HA, Von Haeseler A, Minh BQ. IQ-TREE: A fast and effective stochastic algorithm for estimating maximum-likelihood phylogenies. Mol Biol Evol 2015; 32: 268–274.

72 Kalyaanamoorthy S, Minh BQ, Wong TKF, Von Haeseler A, Jermiin LS. ModelFinder: Fast model selection for accurate phylogenetic estimates. Nat Methods 2017. doi:10.1038/nmeth.4285.

73 Price MN, Dehal PS, Arkin AP. FastTree 2 - Approximately maximum-likelihood trees for large alignments. PLoS One 2010; 5: e9490.

74 Marçais G, Delcher AL, Phillippy AM, Coston R, Salzberg SL, Zimin A. MUMmer4: A fast and versatile genome alignment system. PLoS Comput Biol 2018. doi:10.1371/journal.pcbi.1005944.

