## Supplemental Figures S1-S18 for "Integrative analysis of the shikonin metabolic network identifies new gene connections and reveals evolutionary insight into shikonin biosynthesis"

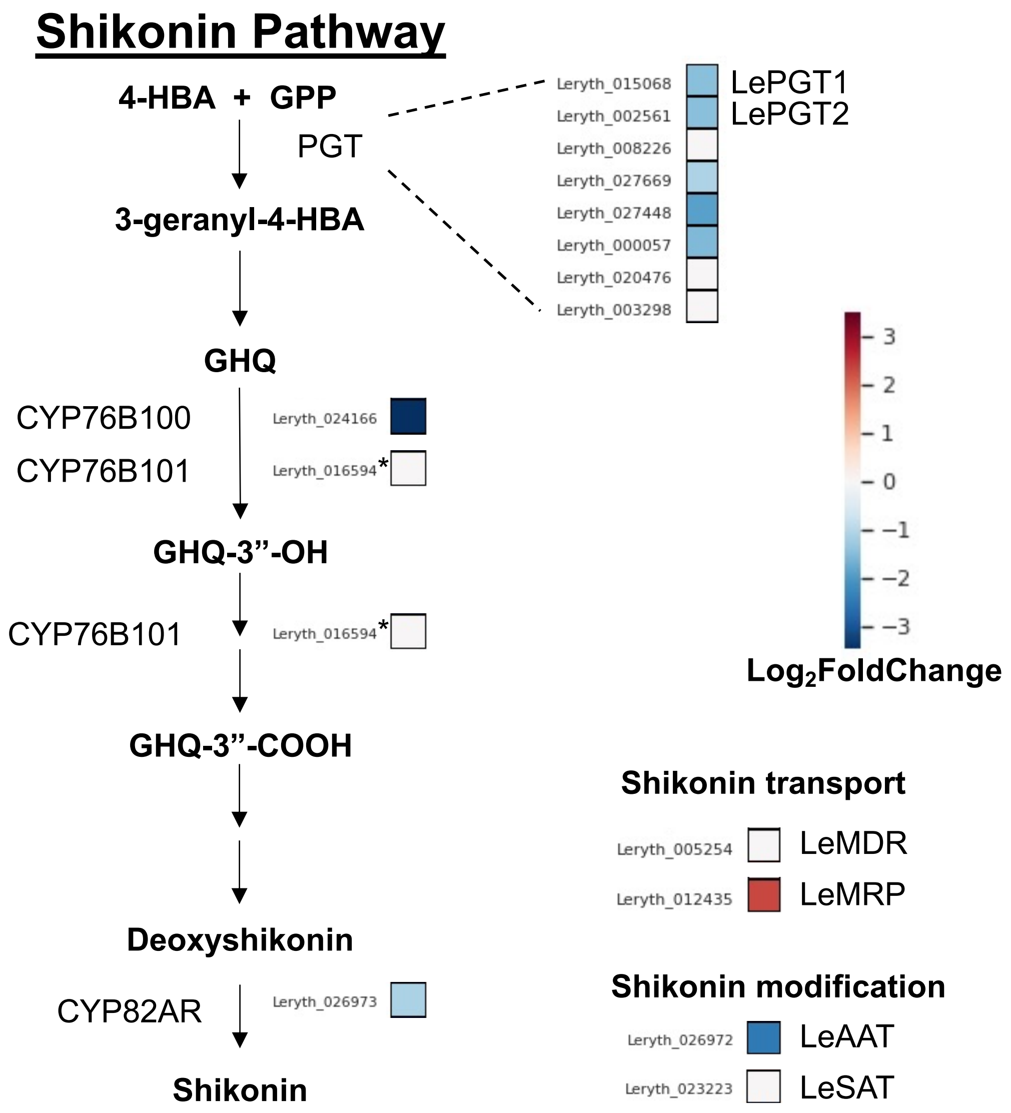


**Fig. S1 Effect of *LeGPPS* RNAi downregulation on expression of shikonin pathway genes.** The average $\log_2$fold-change in expression for each gene in *LeGPPSi*-45 lines compared to *EV-26* lines. The CYP76B101 gene (*) was not included in this DE analysis as the non stranded RNAseq library was unable to distinguish between CYP76B101 (Leryth_016594) and Leryth_016593, which occupies the same genomic location but is encoded on the opposite strand. See Fig. 1 legend for abbreviations and Table S4 for gene descriptions.

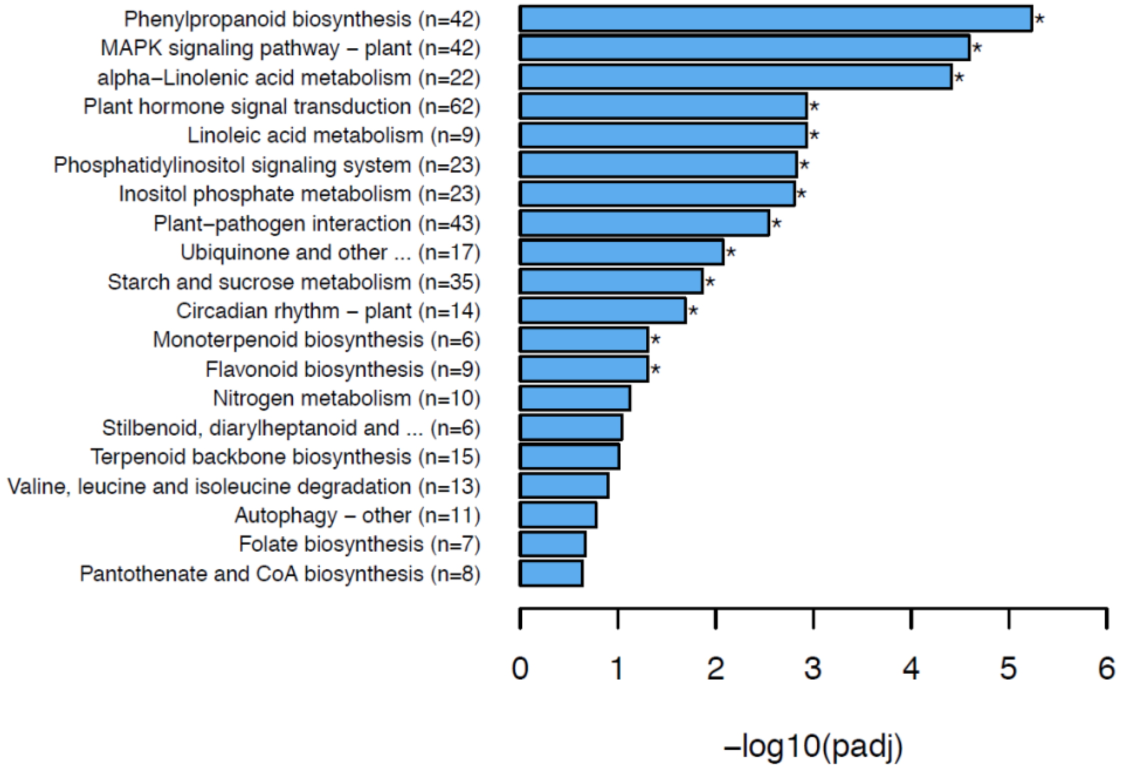


**Fig. S2 Kyoto Encyclopedia of Genes and Genomes (KEGG) term enrichment analysis of genes downregulated in *LeGPPSi*-45 compared to *EV*-26 hairy root lines.** KEGG pathways with corrected p-value < 0.05 were considered significantly enriched by differential expressed genes. *EV*-26, empty-vector control line 26; *LeGPPSi*-45, *LeGPPS*-RNAi line 45.

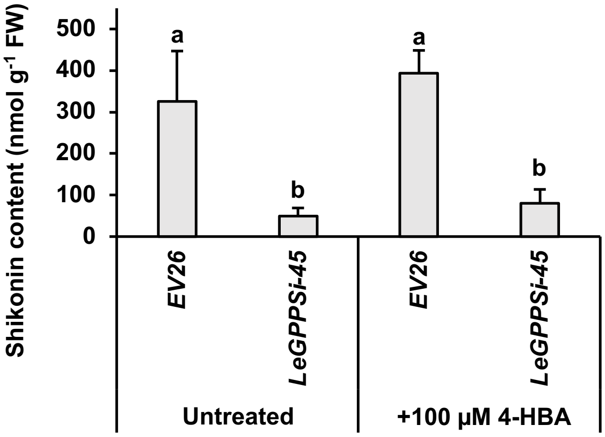


**Fig. S3 Exogenous application of 4-hydroxybenzoate (4-HBA) does not restore shikonin production in *LeGPPSi*-45 lines.** All data are means ± SEM (n = 4 biological replicates). Different letters indicate significant differences via analysis of variance (ANOVA) followed by post-hoc Tukey test (α= 0.05).

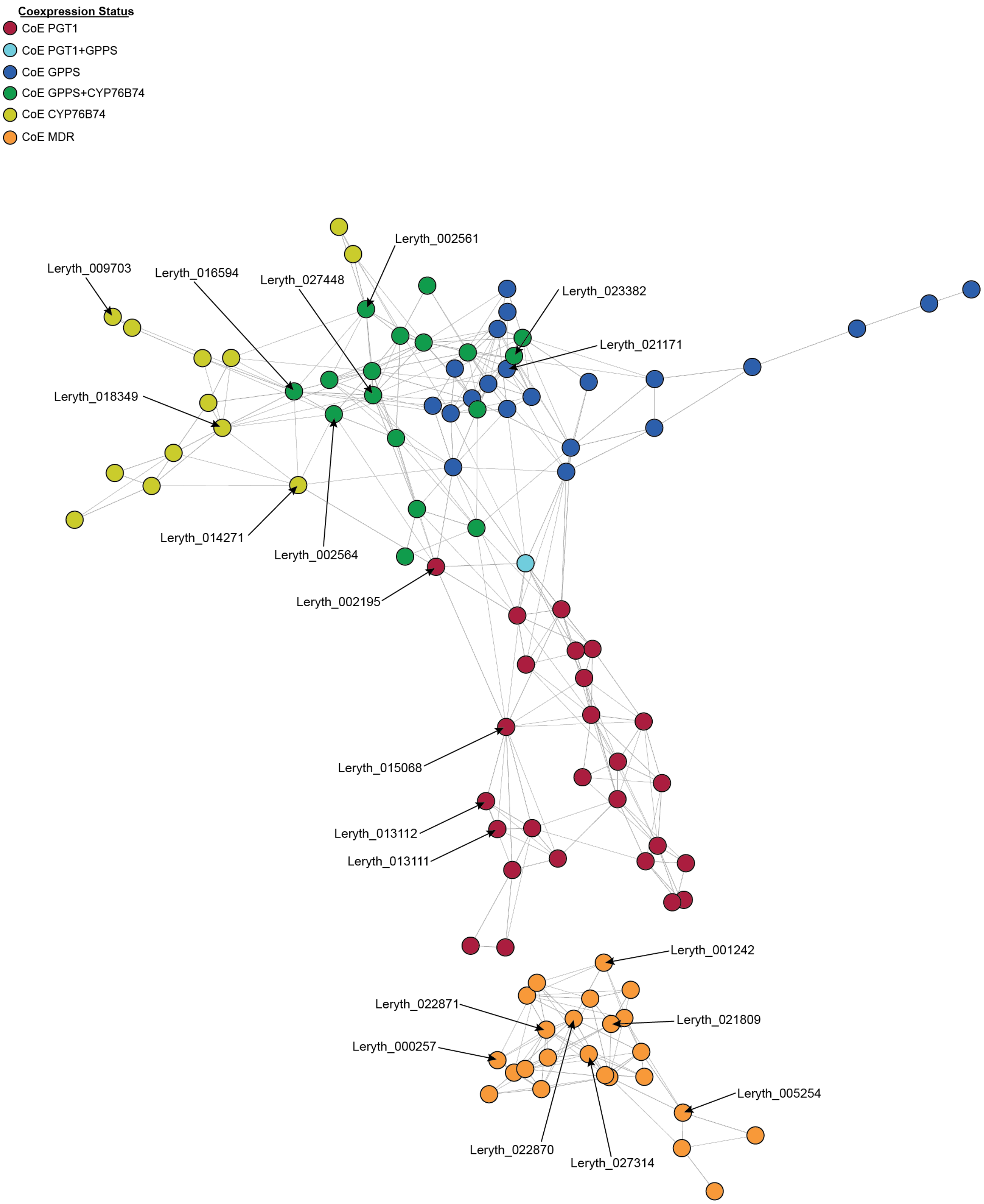


**Fig. S4a Shikonin subnetwork N1.** Network map of genes coexpressed with *LePGT*, *LeGPPS*, *LeCYP76B101*, and *LeMDR* using the N1 global coexpression network. Nodes are colored according to the gene's coexpression status with known shikonin genes. Network maps were drawn using a Fruchterman-Reingold force-directed layout using the edge-weighted spring embedded layout in cytoscape (https://cytoscape.org).

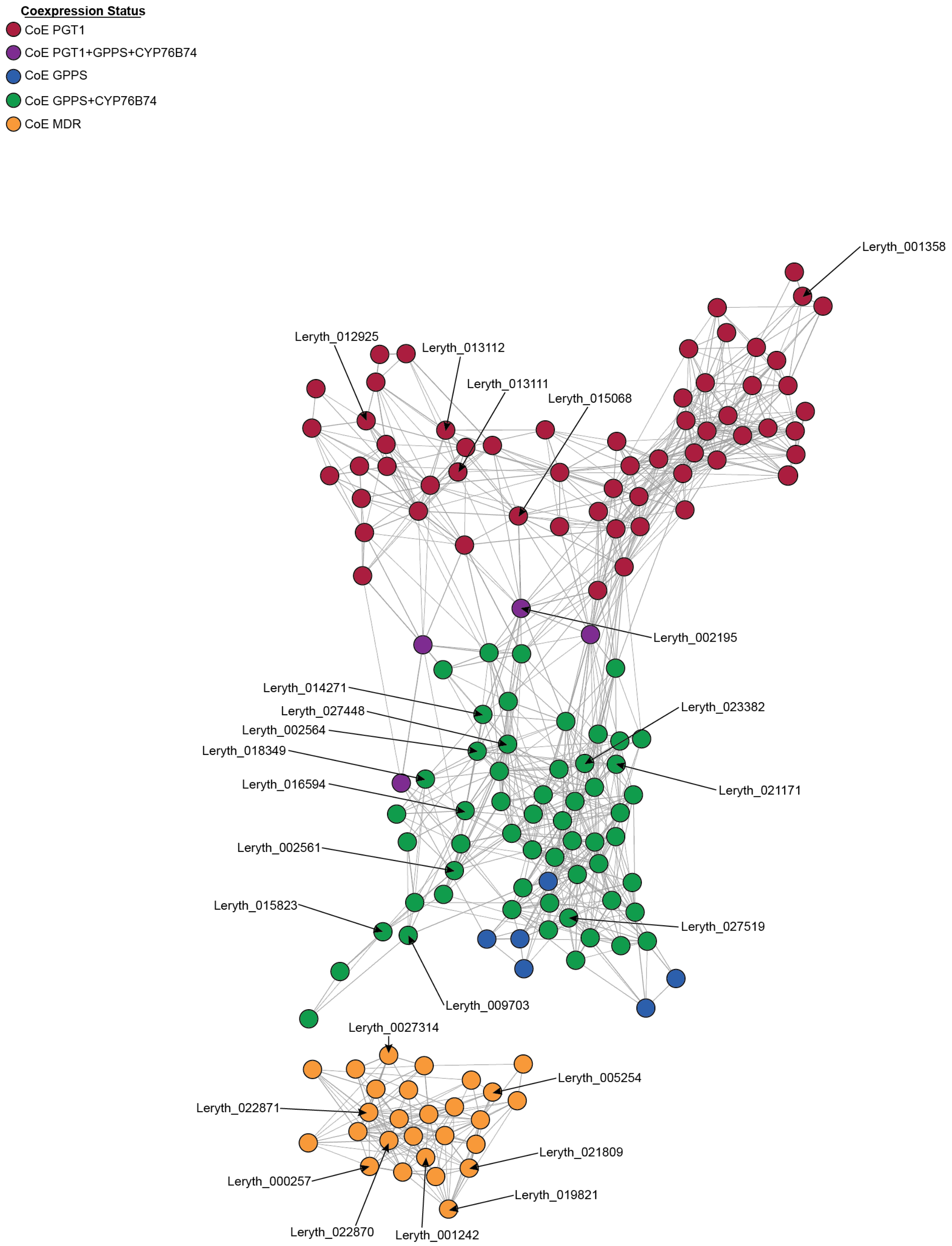


**Fig. S4b Shikonin subnetwork N2.** Network map of genes coexpressed with *LePGT*, *LeGPPS*, *LeCYP76B101*, and *LeMDR* using the N2 global coexpression network. Network maps are drawn as described in Supplemental Figure S1a.

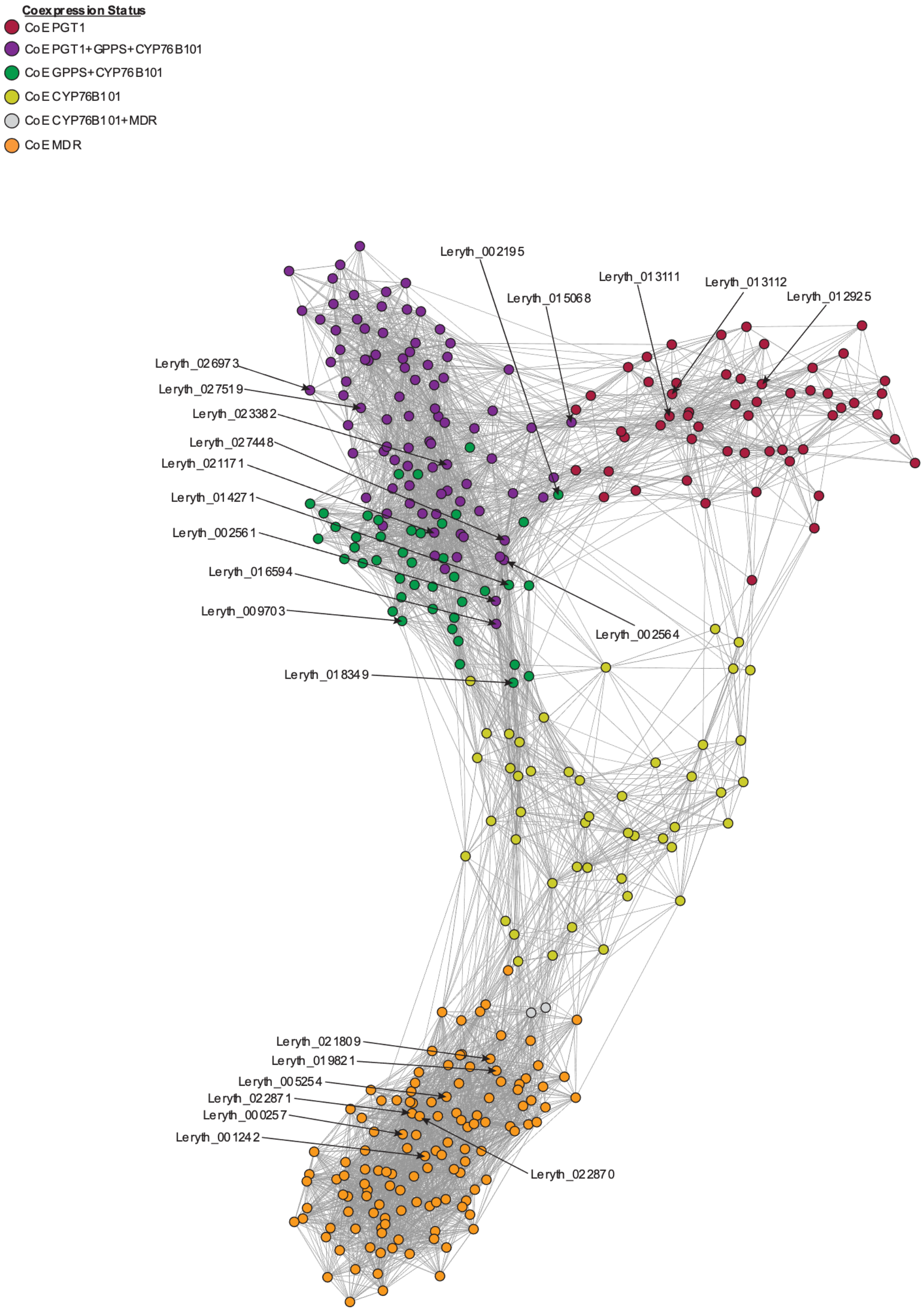


**Fig. S4c Shikonin subnetwork N3.** Network map of genes coexpressed with *LePGT*, *LeGPPS*, *LeCYP76B101*, and *LeMDR* using the N3 global coexpression network. Network maps are drawn as described in Supplemental Figure S1a.

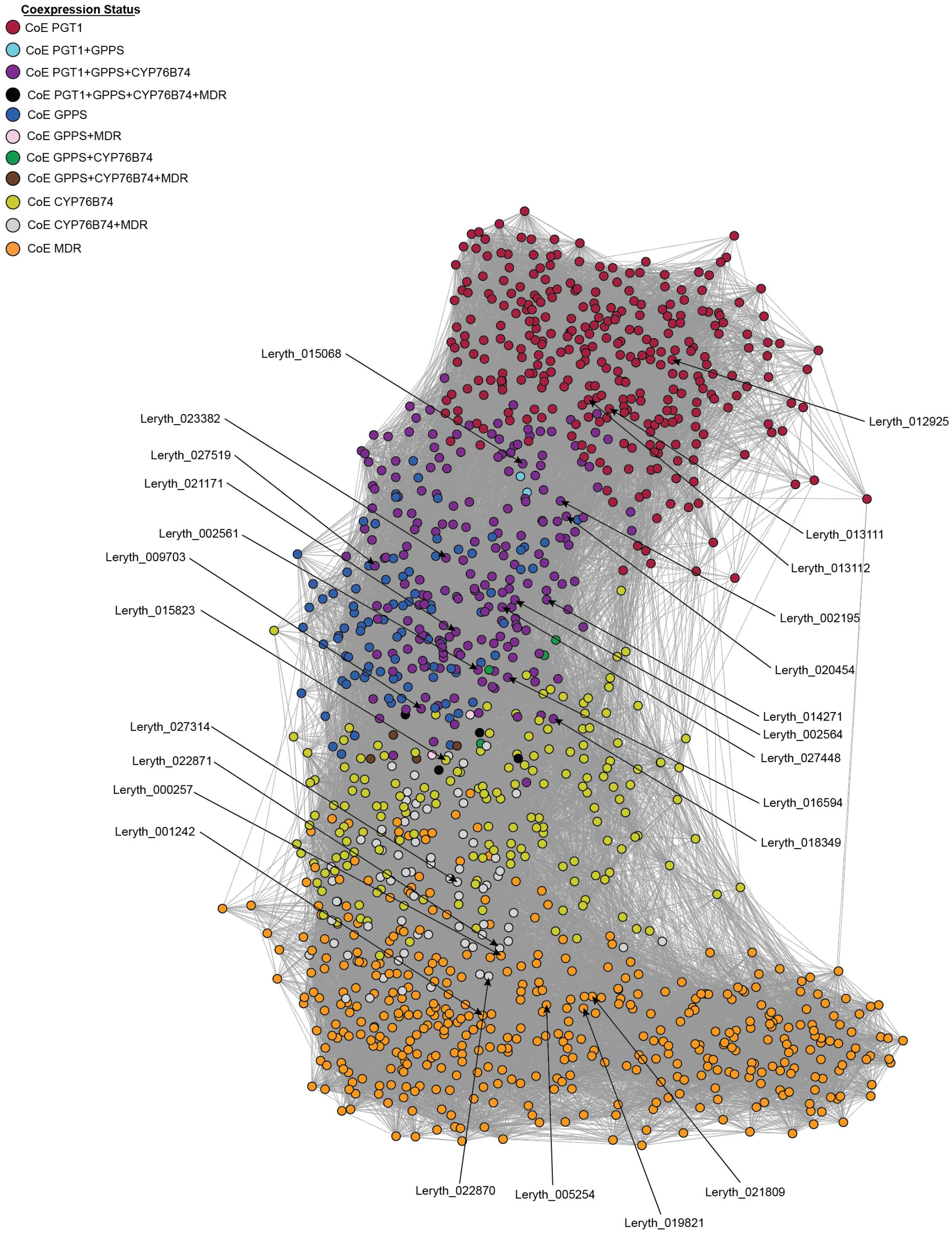


**Fig. S4d Shikonin subnetwork N4.** Network map of genes coexpressed with *LePGT*, *LeGPPS*, *LeCYP76B101*, and *LeMDR* using the N4 global coexpression network. Network maps are drawn as described in Supplemental Figure S1a.

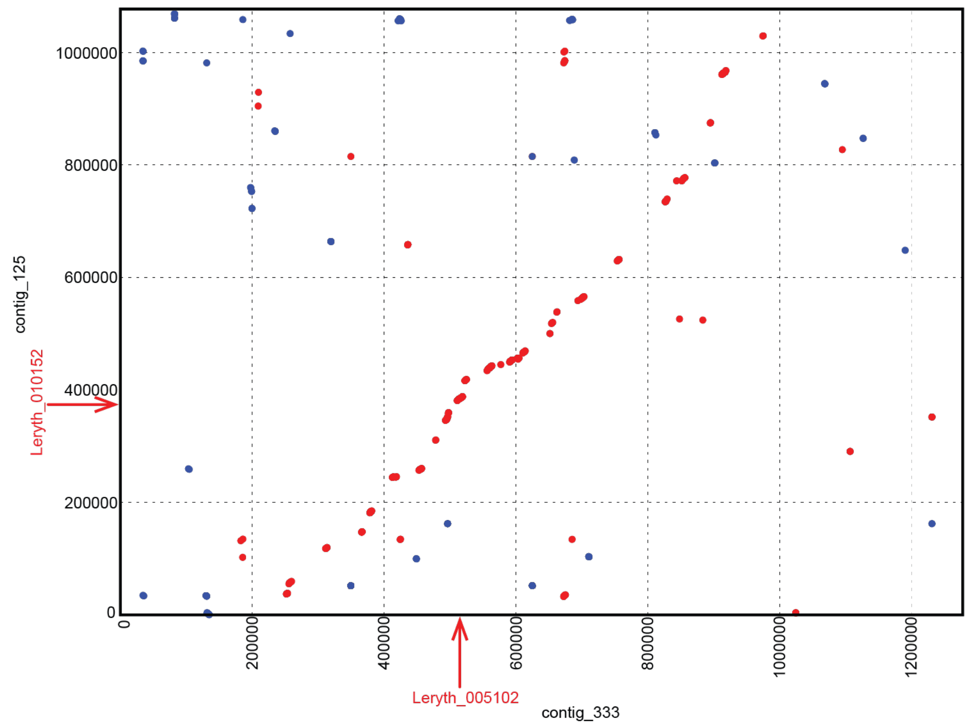


**Fig. S5 Shared synteny between FPPS homologs**. Mummerplot of syntenic block containing farnesyl pyrophosphate synthase homologs Leryth_005102 (*LeFPPS1*) and Leryth_010152 *(LeFPPS3*). The locations of the homologs are indicated by the red arrows.

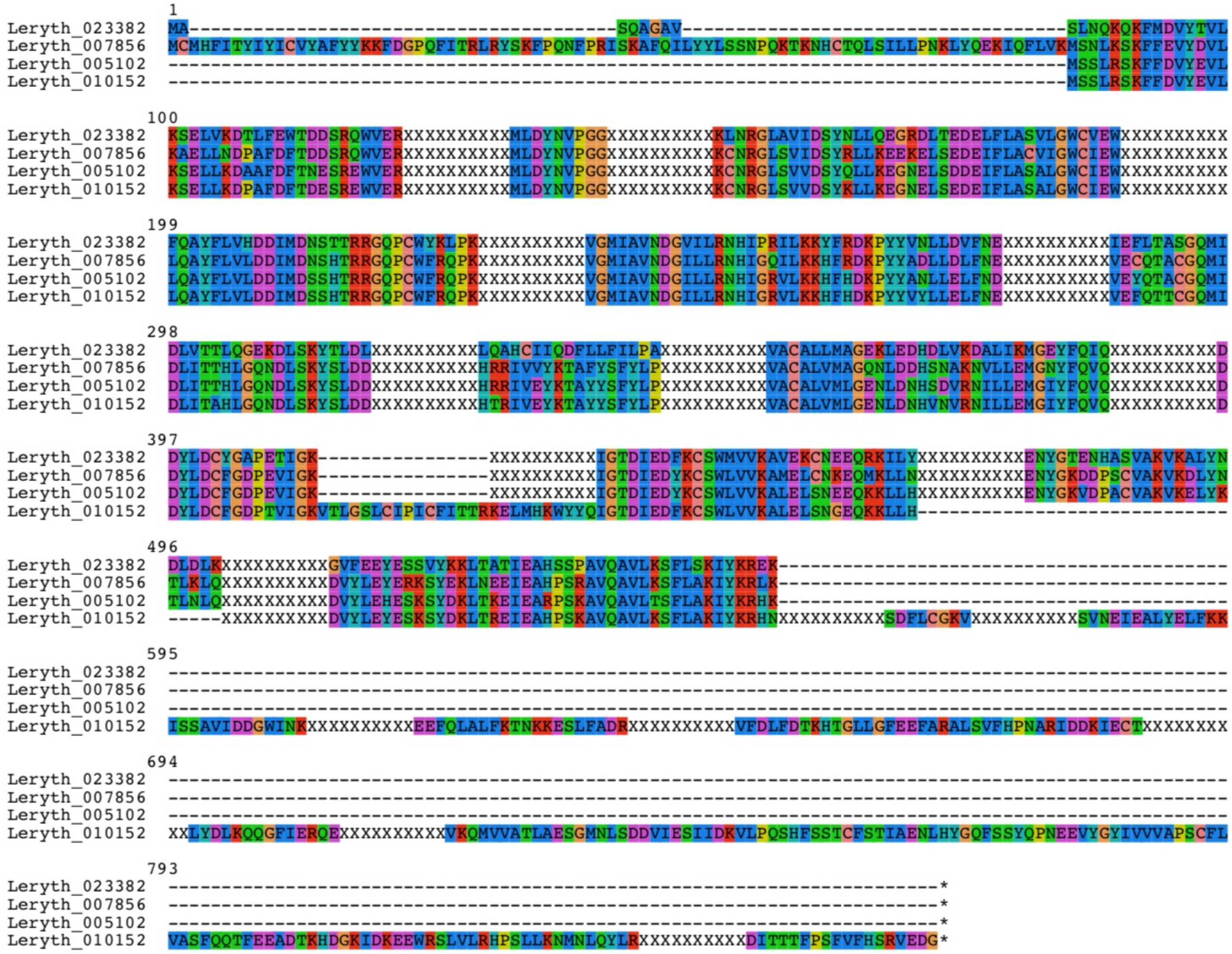


**Fig. S6 Conservation of intron locations in the *FPPS* gene family**. Multiple sequence alignment showing conservation of intron locations (indicated by xxxxxxxxxx) between LeGPPS (Leryth_023382), LeFPPS1 (Leryth_005102), LeFPPS2 (Leryth_007856), and LeFPPS3 (Leryth_010152).

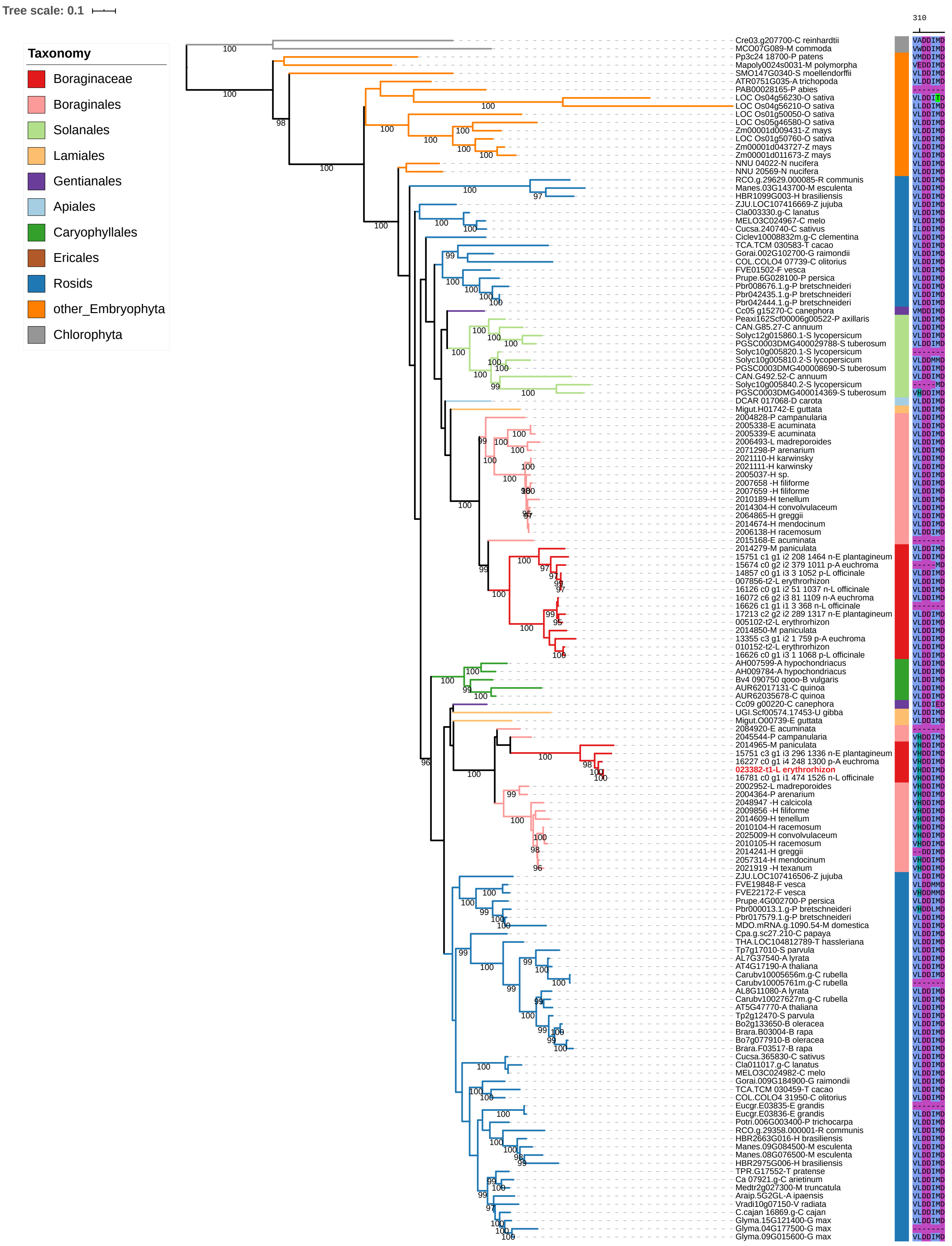


**Fig S7 Maximum likelihood phylogeny of FPPS gene family in green plants.** Nodes with IQ-TREE support values > 95 are indicated by numbers on the preceding branch. The branches and outer color bar are color-coded to match the taxonomic classification of each sequence. The tree is rooted on Chlorophytes. Non-Boraginales sequences containing a Histidine residue adjacent to the conserved Asp-rich motif are indicated by an asterisk (Fv = F. vesca, Pb = P. bretschneideri, St = S. tuberosum).

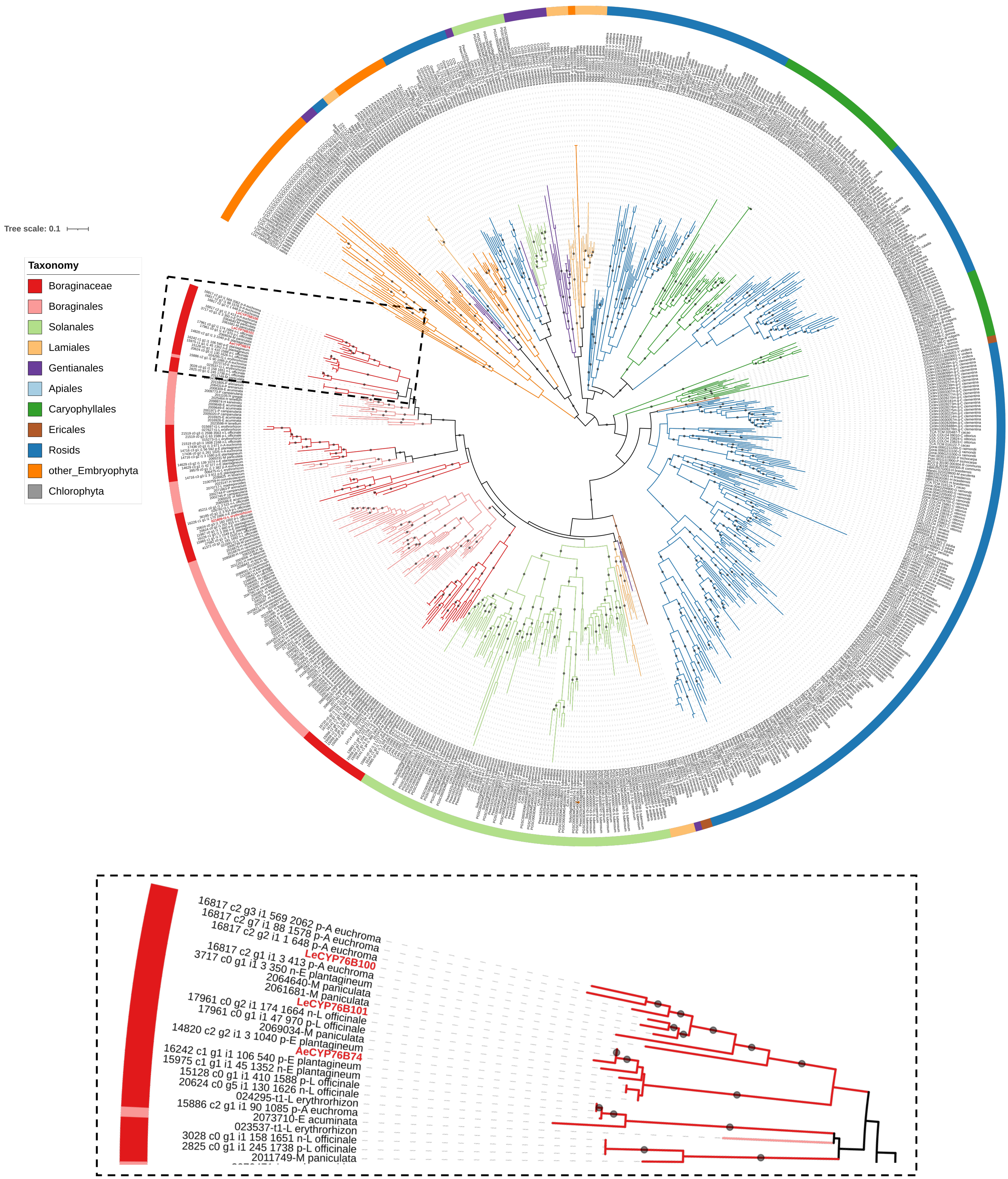


**Figure S8 Maximum likelihood phylogeny of cytochrome CYP76B gene family in green plants.** Nodes with IQ-TREE support values > 95 are indicated by grey circles on the preceding branch. The branches and outer color bar are color-coded to match the taxonomic classification of each sequence. The tree is rooted based on rough guide tree of entire cytochrome P450 gene family (Plaza Dicots 4.0 HOM04D000003).

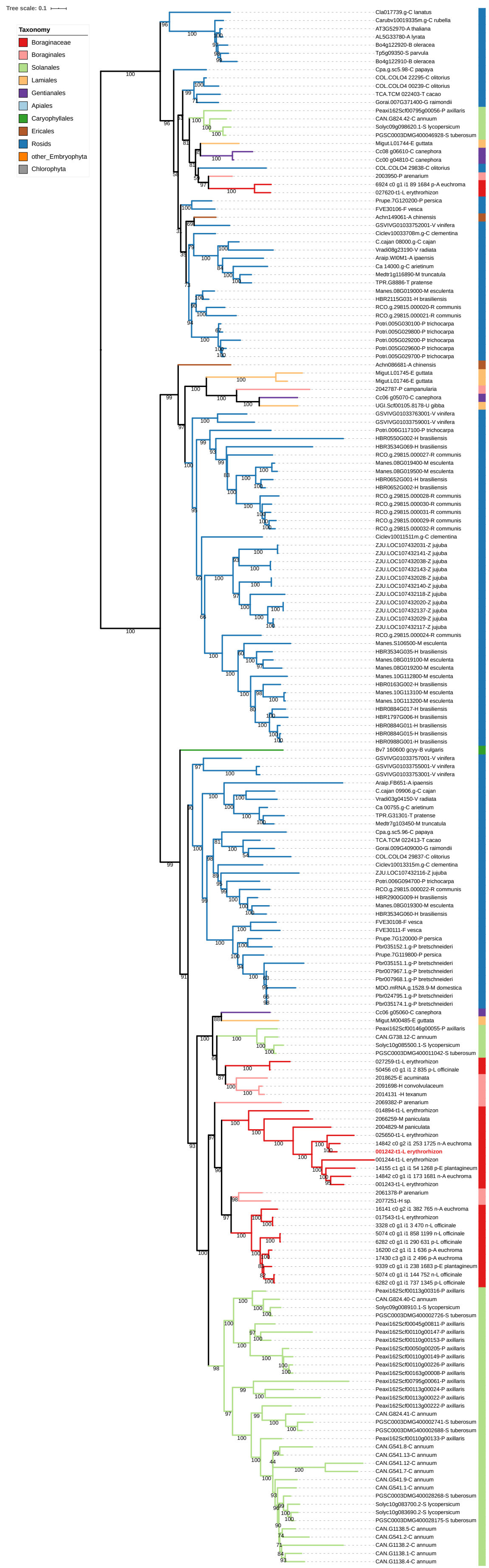


**Figure S9 Maximum likelihood phylogeny of cytochrome P450 candidate Leryth_001242 and homologs in green plants.** Nodes with IQ-TREE support values > 95 are indicated by numbers on the preceding branch. The branches and outer color bar are color-coded to match the taxonomic classification of each sequence. The tree is rooted based on rough guide tree of entire cytochrome P450 gene family (Plaza Dicots 4.0 HOM04D000003).

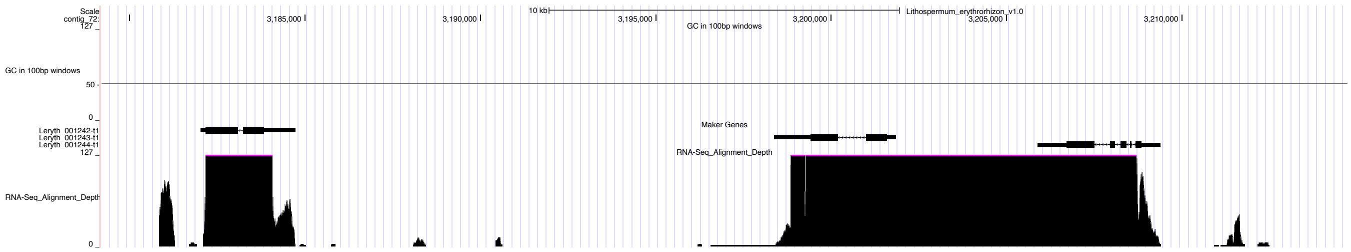


**Figure S10 UCSC Genome Browser region for cytochrome P450 candidate Leryth_001242.**

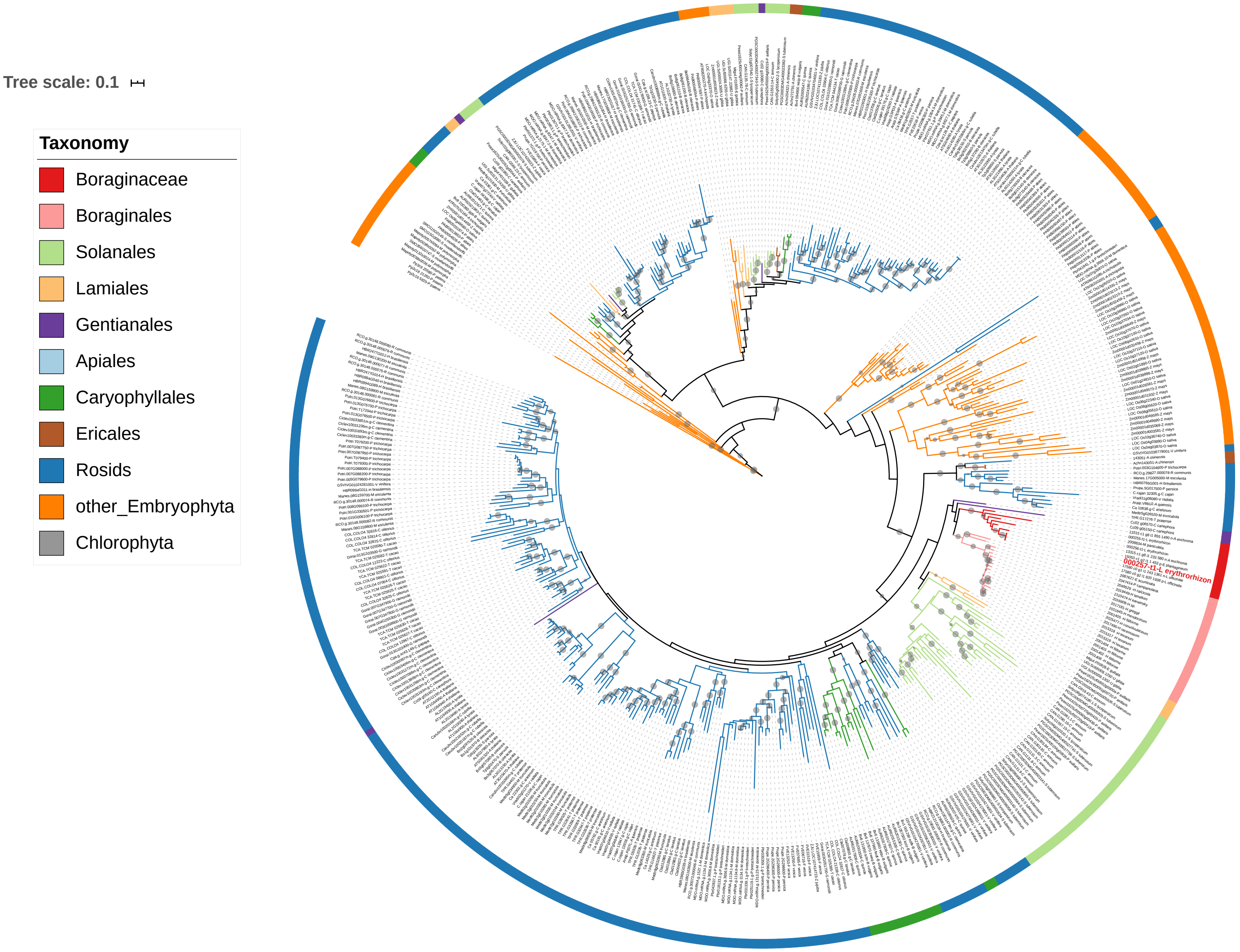


**Figure S11 Maximum likelihood phylogeny of cytochrome P450 candidate Leryth_000257 and homologs in green plants.** Nodes with IQ-TREE support values > 95 are indicated by grey circles on the preceding branch. The branches and outer color bar are color-coded to match the taxonomic classification of each sequence. Tree is midpoint rooted. The tree is rooted based on rough guide tree of entire cytochrome P450 gene family (Plaza Dicots 4.0 HOM04D000435)..

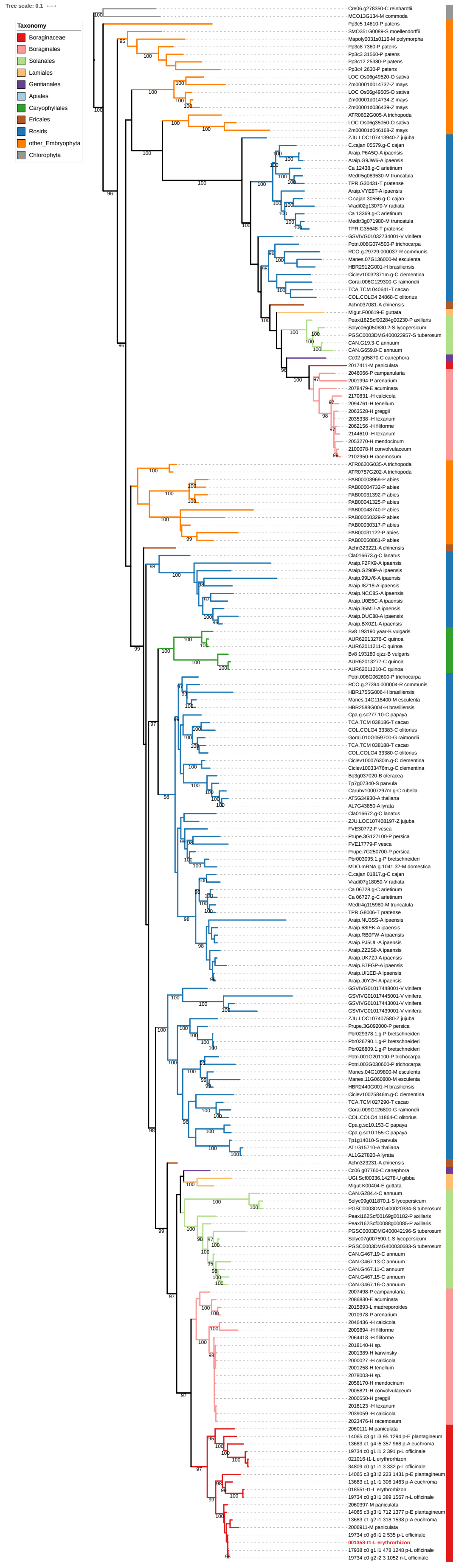


**Figure S12 Maximum likelihood phylogeny of the prephenate dehydrogenase gene family in green plants.** Nodes with IQ-TREE support values > 95 are indicated by numbers on the preceding branch. The branches and outer color bar are color-coded to match the taxonomic classification of each sequence. The tree is rooted on Chlorophytes.

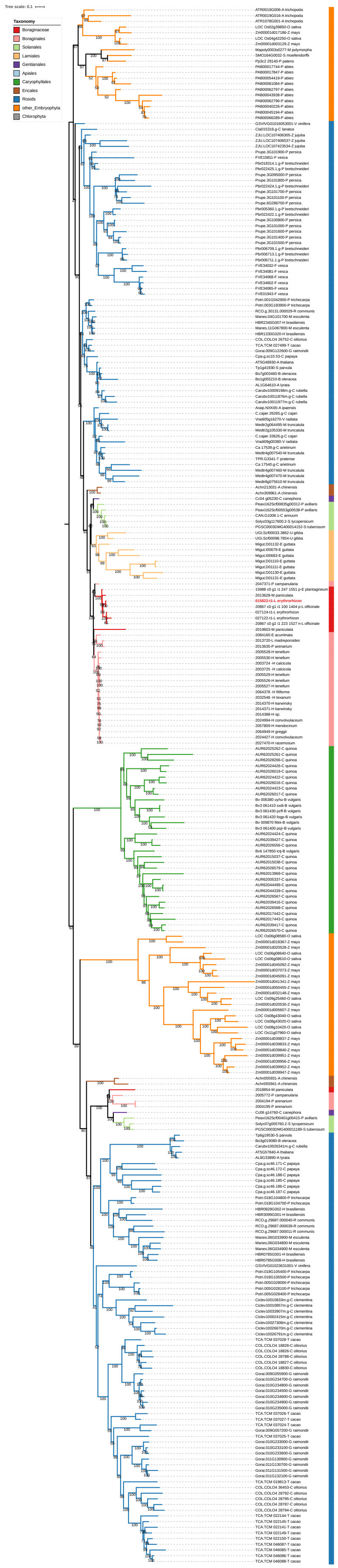


**Figure S13 Maximum likelihood phylogeny of transferase candidate Leryth_015823 and homologs in green plants.** Nodes with IQ-TREE support values > 95 are indicated by numbers on the preceding branch. The branches and outer color bar are color-coded to match the taxonomic classification of each sequence. The tree is rooted based on rough guide tree of entire AT gene family (Plaza Dicots 4.0 HOM04D000075).

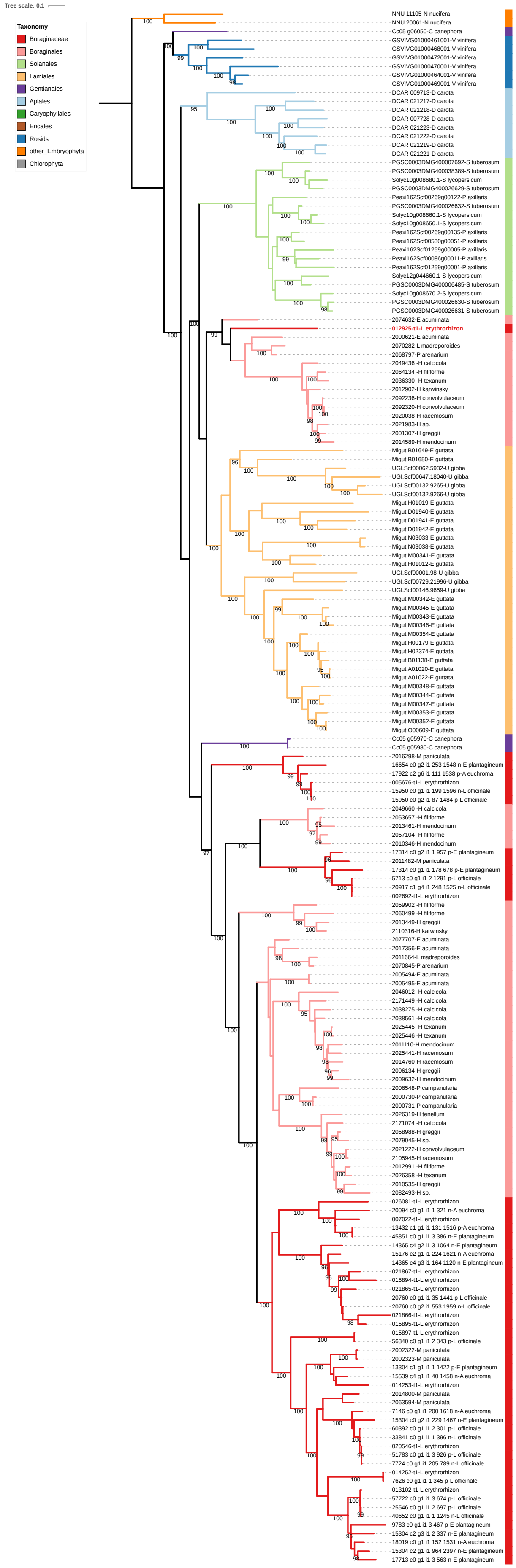


**Figure S14 Maximum likelihood phylogeny of transferase Candidate Leryth_012925 and homologs in green plants.** Nodes with IQ-TREE support values > 95 are indicated by numbers on the preceding branch. The branches and outer color bar are color-coded to match the taxonomic classification of each sequence. The tree is rooted based on rough guide tree of entire AT gene family (Plaza Dicots 4.0 HOM04D000339).

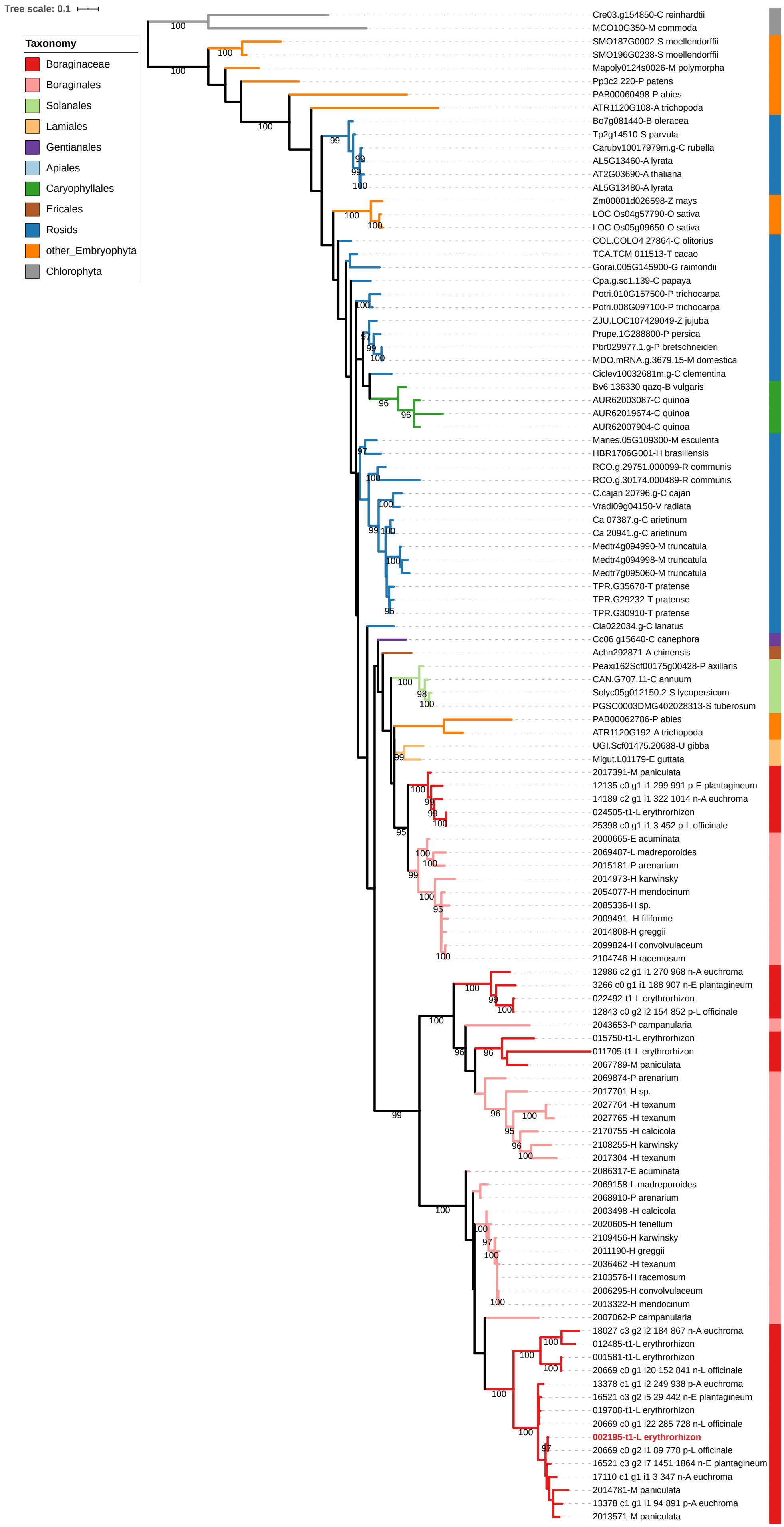


**Figure S15 Maximum likelihood phylogeny of COQ4 gene family in green plants.** Nodes with IQ-TREE support values > 95 are indicated by numbers on the preceding branch. The branches and outer color bar are color-coded to match the taxonomic classification of each sequence. The tree is rooted on Chlorophytes.

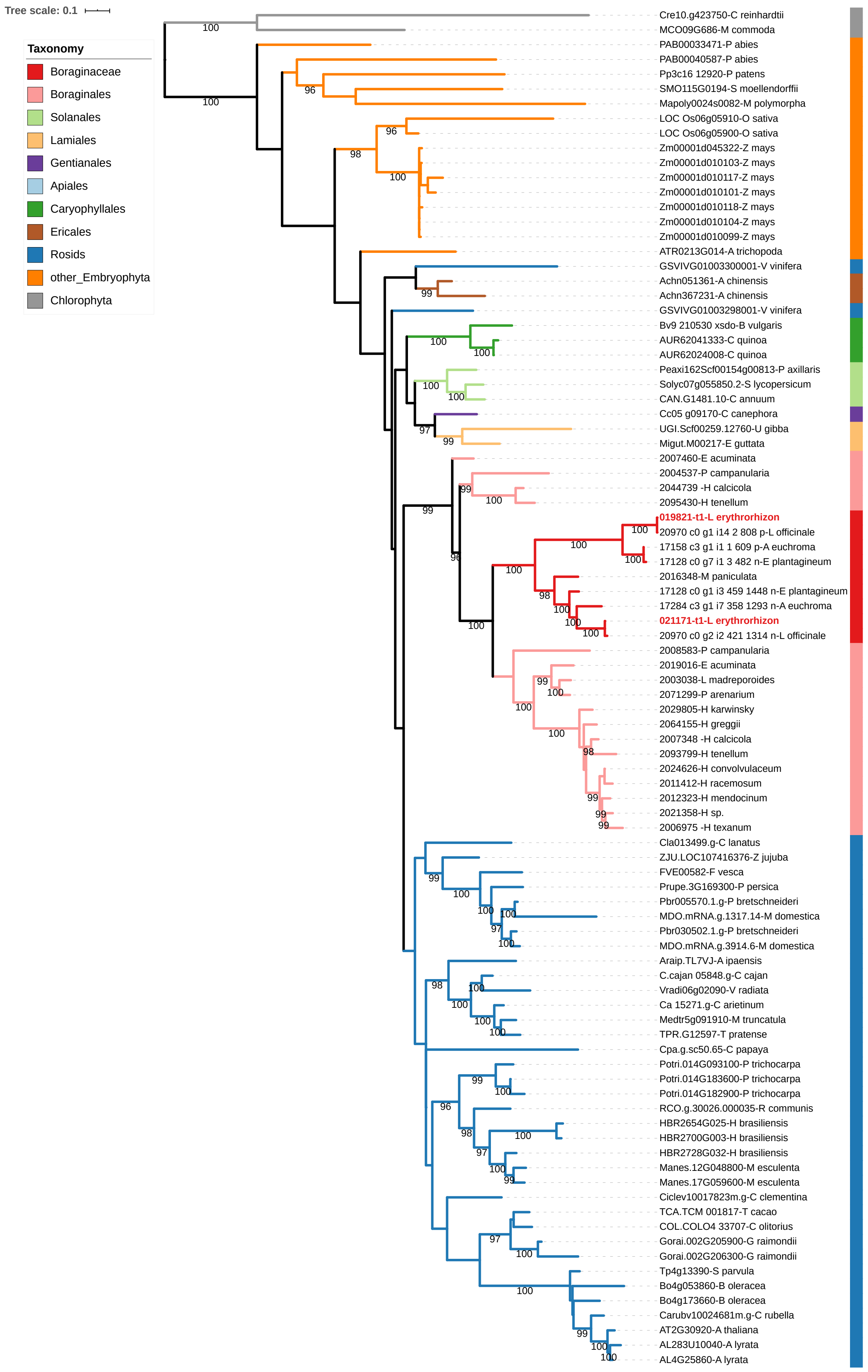


**Figure S16 Maximum likelihood phylogeny of the COQ3 gene family in green plants.** Nodes with IQ-TREE support values > 95 are indicated by numbers on the preceding branch. The branches and outer color bar are color-coded to match the taxonomic classification of each sequence. The tree is rooted on Chlorophytes.

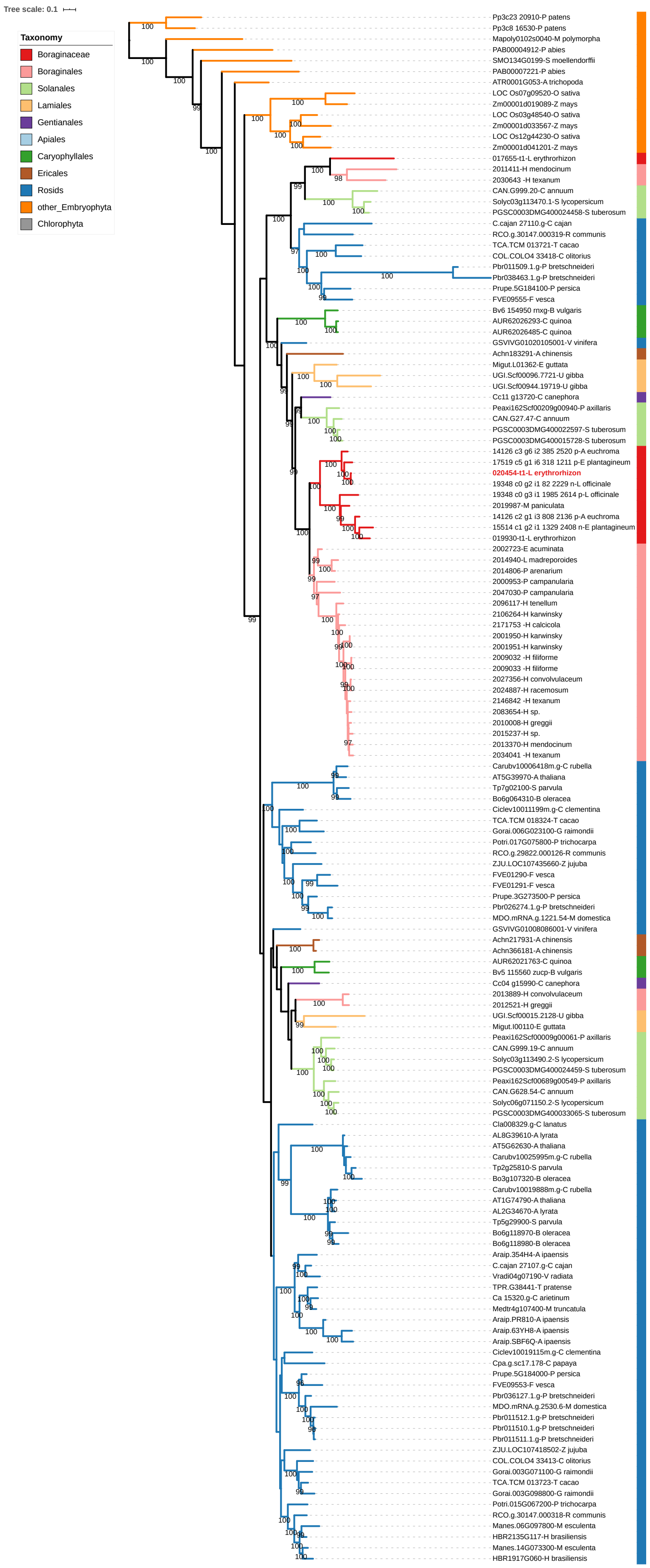


**Figure S17 Maximum likelihood phylogeny of quinoprotein dehydrogenase gene family in green plants.** Nodes with IQ-TREE support values > 95 are indicated by numbers on the preceding branch. The branches and outer color bar are color-coded to match the taxonomic classification of each sequence. The tree is rooted on mosses.

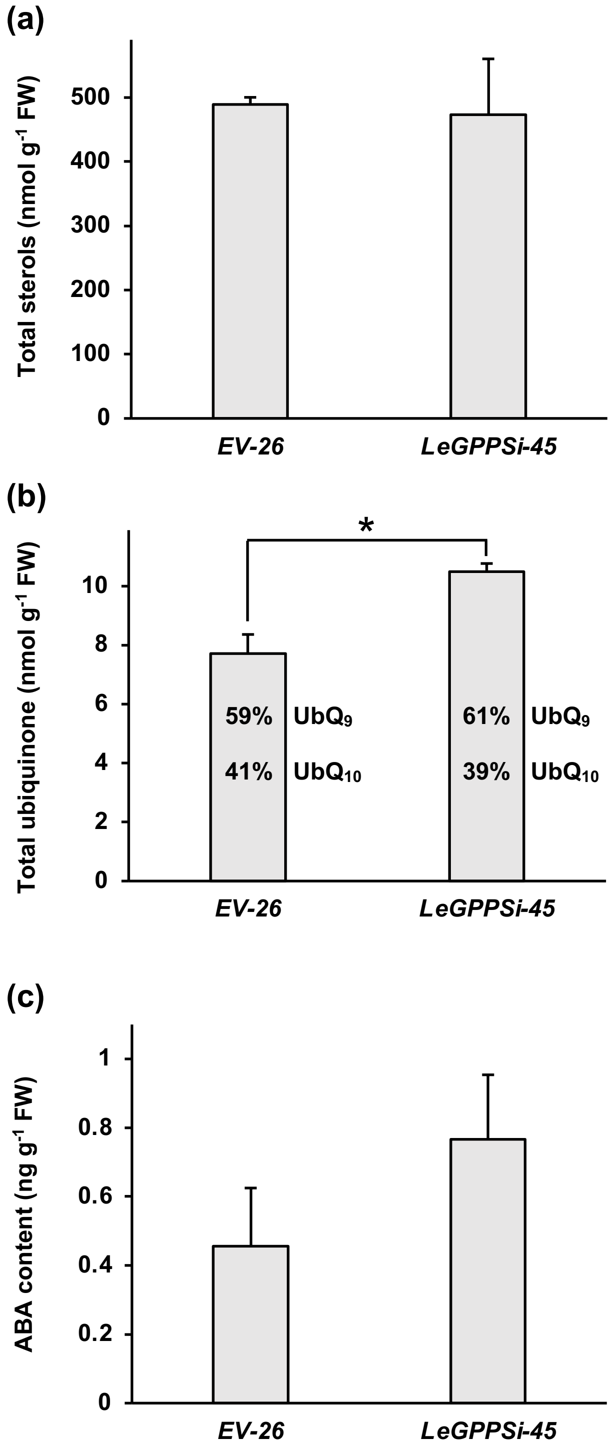


**Figure S18 Pool sizes of sterols (a), ubiquinones (b), and abscisic acid (ABA) (c) measured in empty-vector control line 26 (*EV*-26) and *LeGPPS* RNAi line 45 (*LeGPPSi*-45).** Metabolites were measure at 6 d after transfer of 14-d-old hairy roots to M9 and darkness. All data are means ± SEM (n = 3–4 biological replicates). Statistically significant differences are indicated (*P < 0.05, Student's *t* test).
